# Unveiling the potential of neuron-targeted dendriplexes for siRNA delivery using a PNS-CNS-on-Chip

**DOI:** 10.1101/2024.09.05.611457

**Authors:** Ana P. Spencer, Miguel Xavier, Sofia C. Guimarães, Adriana Vilaça, Ariel Ionescu, Rafael Santos, María Lázaro, Eran Perlson, Victoria Leiro, Ben M. Maoz, Ana P. Pêgo

## Abstract

Neurological disorders, a leading global cause of death, encompass conditions affecting the peripheral and central nervous systems (PNS and CNS, respectively). Limited axon regeneration is a significant challenge in these disorders, and it is linked to proteins like PTEN. RNA-based therapeutics, particularly siRNAs, hold potential for silencing these inhibitory pathways, but their clinical application is hindered by poor stability and cellular uptake. Our study addressed this challenge with the development of novel, fully biodegradable dendritic nanoparticles designed specifically for neuron targeting. These nanoparticles were functionalized with the neurotropic binding domain of tetanus toxin, enhancing selective neuronal targeting and cellular internalization. We demonstrated that these dendriplexes not only maintain biocompatibility and efficient siRNA delivery in neuronal cultures but also significantly enhance axonal growth, as shown in microfluidic models. In a groundbreaking PNS-CNS-on-Chip, dendriplexes exhibited effective migration from PNS to CNS neurons, highlighting their potential for targeted therapeutic delivery. This study pioneers the application of microfluidics to demonstrate the CNS targeting of dendriplexes, paving the way for innovative treatments in the field of nanomedicine.

## Introduction

Considered the second leading cause of death and responsible for nearly 10 million deaths annually, with a progressively aging population, neurological disorders have also emerged as the foremost contributor to age-associated functional and cognitive limitations worldwide [1, 2]. These conditions include those affecting the peripheral nervous system (PNS), resulting in altered sensory and motor capabilities, namely diabetic neuropathy [3], Guillain-Barré syndrome [4], and Charcot-Marie-Tooth disease [5] in which patients often experience symptoms like numbness, tingling, muscle weakness, and pain. The central nervous system (CNS) is also vulnerable to various disorders, including Alzheimer’s [6], Parkinson’s [7], and Huntington’s [8], examples of the vast panoply of neurodegenerative diseases. Autoimmune disorders like multiple sclerosis, where the immune system mistakenly targets the myelin sheath surrounding axons, causing disruptions in nerve conduction, also afflict the CNS with severe consequences to patients. Additionally, stroke [9], traumatic brain injury [10], and brain tumours [11] can also cause severe disruptions to the CNS.

A critical challenge across many of these conditions is the limited capacity for axon regeneration, which is often attributed to the expression of inhibitory proteins like phosphatase and tensin homolog (PTEN) [12–14]. PTEN’s suppression of axonal growth and promotion of growth cone collapse severely hinder neural repair, making it a crucial target for therapeutic intervention [13]. In fact, by deactivating or inhibiting PTEN, either through genetic manipulation, using DNA- or RNA-based therapeutics, or pharmaceutical interventions, enhanced axon regrowth and improved neuronal regeneration were observed [12].

The use of DNA- or RNA-based therapeutics is evolving dramatically, with recent groundbreaking research on mRNA earning the 2023 Nobel Prize in Medicine [15]. NA-based therapeutics, particularly siRNAs, offer a precise approach to gene silencing but are plagued by limitations, including rapid degradation, inefficient cellular uptake, and off-target effects. This is precisely where nanotechnology emerges as an essential solution. Nanoparticles (NPs) created by integrating delivery vectors and nucleic acids represent a remarkable fusion of science and technology. These NPs can be engineered to carry and protect RNA molecules, thus significantly enhancing their stability and internalization by cells [16]. One of the most significant advantages of this nanotechnological innovation lies in the precision with which the surface of NPs can be strategically customized using a diverse array of ligands, including targeting moieties for specific cell populations. The targeting capability is crucial for improving the efficiency of delivering therapeutic payloads while minimizing the risk of side effects [17].

Among the delivery vectors capable of delivering nucleic acids, dendrimers have emerged as promising candidates and may represent the long-awaited nanomedicine solution to treat and reduce the burden of various diseases [18–23]. Renowned as unique nanocarriers, dendrimers boast a well-defined and tri-dimensional architecture, consisting of concentric layers offering precise control over size [24, 25]. Their peripheral layers allow customization with diverse surface functionalities, leading to a versatile and distinctive structure. This makes dendrimers attractive for various applications with the further possibility of targeted ligand attachment to their structure or their corresponding NPs [18].

However, existing NP systems often face challenges such as suboptimal biocompatibility, low transfection efficiency, limited target specificity, and potential toxicity. Many conventional NPs fail to traverse complex neural interfaces, like the blood-brain barrier, limiting their therapeutic impact on CNS disorders. Furthermore, commonly used NPs, such as non-degradable dendrimers, frequently exhibit poor biodegradability, raising concerns about long-term toxicity and clearance [26, 27].

To address these limitations, we have designed and developed a novel family of fully biodegradable PEG-GATGE (poly(Ethylene Glycol)-Gallic Acid-Triethylene Glycol Ester) dendritic block copolymers [19, 28], engineered to improve the delivery of siRNAs specifically to neuronal cells. To act as siRNA delivery vectors, the periphery of our dendritic block copolymers was functionalized with benzylamine groups (fbB dendrimer) [19, 28]. Unlike commonly used dendrimers, these vectors completely degrade under physiological conditions. Furthermore, fbB dendrimers have shown an impressive capacity to effectively bind and safeguard siRNA in highly compact and small dendriplexes (siRNA-dendrimer complexes), measuring less than 60 nm. Also, this fully biodegradable vector has displayed a low cytotoxic profile [19, 28] with efficient intracellular siRNA release, resulting in significant gene downregulation in different cell lines [19].

To exploit the use of these dendriplexes for neuronal treatment, we have developed a new, safe, and efficient neuron-targeted (Tg) nanocarrier based on fbB dendrimers and siRNA designed to downregulate PTEN expression (siPTEN). To attain neurotropism, our dendriplexes were functionalized with the non-toxic C-terminal fragment of the tetanus neurotoxin (TeNT) heavy chain (HC) [29]. In prior studies, we explored trimethyl chitosan (TMC) NPs that were functionalized with the HC fragment to specifically target PNS neurons and undergo retrograde transportation to the cell body [30, 31]. However, HC’s full potential in CNS-targeting and transcytosis remained unexplored.

To explore this, three advanced microfluidic platforms were developed to explore the features of our fully biodegradable and Tg dendritic NPs, focusing specifically on intra-neuronal migration, biological effects, and transcytosis between neurons. For this, it was crucial to create a novel microfluidic-based smart model, the PNS-CNS-on-Chip, integrating microelectrode arrays (MEAs), and engineered to simulate various aspects of the complex PNS-CNS interface, allowing the validation of the targeted delivery capabilities of our NPs.

By exploring nanotechnological tools and microfluidics, this study addressed the critical need for targeted, efficient, and biocompatible delivery systems in neurodegenerative disease treatment. We have demonstrated that our HC-tethered dendriplexes could not only target neurons from both the PNS and CNS but also showed, for the first time, that NPs can reach the CNS and actuate centrally following a minimally invasive and clinically significant peripheral administration route.

## 2. Materials and methods

### 2.1. Materials and animals

The synthesis of generation 3 of the PEG-GATGE dendritic block copolymer, along with its peripheral functionalization by “click” chemistry with alkynated benzylaminium salt (4-ethynyl-benzenemethanamine·HCl), and its characterization was conducted according to our previously reported procedure [19]. Double-stranded RNAs targeting enhanced green fluorescence protein (GFP) or PTEN (designated as siGFP and siPTEN, respectively) were used for the dendriplexes’ preparation [28]. A mimetic of siGFP (siRNAmi) labelled at the 5′ end of the sense strand with cyanine 5 (Cy5) was used when it was intended to monitor the dendriplexes intracellular path. All sequences (detailed in Table S1, Support Information – SI) were supplied by Integrated DNA Technologies (USA).

All animal experiments were conducted following European Union guidelines (EU Directive 2010/63/EU) and Portuguese law (DL 113/2013), with the utmost concern to minimize animal suffering as well as the number of animals used. All the procedures with animals were previously approved by the Portuguese official authority on animal welfare and experimentation (Direção Geral de Alimentação e Veterinária - DGAV), Instituto de Investigação e Inovação em Saúde (i3S) Ethics Committee (CEA, i3S) and Animal Ethics Committee of Tel Aviv University (TAU). Animals were kept in an enriched housing environment with *ad libitum* feed and water supply. Furthermore, animals were kept under a 12/12-hour light/dark cycle with controlled ambient temperature and humidity either at the i3S or TAU animal facilities. Mouse embryos (E12.5 and E16.5) from pregnant 2–3-month-old female C57BL/6 mice were used to obtain the motor and cortical neuron cultures. For downregulation studies, HB9::GFP mice were used. These were obtained from the Jackson Laboratories, and the colony was maintained by breeding with ICR mice. Following ethical guidelines and institutional protocols, euthanasia of the mice was conducted using humane and approved methods. Firstly, mice were placed in a CO_2_ chamber, where they were exposed to carbon dioxide until unconsciousness was achieved. Following CO_2_ exposure, cervical dislocation was performed to ensure rapid and effective euthanasia.

### 2.2. HC fragment production and purification

The plasmid encoding the HC fragment was generously provided by Prof. Neil Fairweather (King’s College, UK) and was produced using a competent BL21 *E. coli* strain and purified as previously described [32].

### 2.3. Surface functionalization of siRNA-dendrimer complexes with TeNT HC fragment

The dendriplexes formed between the fbB dendritic block copolymer and siRNA (siGFP, siPTEN, or Cy5-siRNAmi) were prepared and characterized according to previously established methods [19, 28]. In summary, dendriplexes were prepared at three different N/P ratios (5, 10 and 20), where N represents the moles of primary amines in the dendritic structure and P is the moles of phosphate groups in the siRNA backbone. Initially, siRNA (20 µM, siRNA final concentration of 0.6 μM) was added to different volumes of fbB dendrimer solution (5.5 mg/mL) (fbB final concentration after filtration of a 6.0 mg/mL fbB solution with 0.45 μm PTFE filter (VWR), final volume 100 μL), in 20 mM HEPES (4-(2-hydroxyethyl)-1 piperazineethanesulfonic acid, Gibco) with 5% (w/v) glucose (Fisher Chemical^TM^) pH 7.4 and nuclease-free water (NF water, Qiagen). Then, dendriplex solutions were vortexed for 10 seconds at medium speed (velocity 3, VV3 vortex, VWR) followed by an incubation of 30 minutes at room temperature (RT). To prepare Tg dendriplexes, dendriplex solutions were incubated with a heterofunctional 1 kDa PEG spacer (Creative PEGWorks) bearing an N-hydroxysuccinimide (NHS) and a maleimide (MAL) end group (0.5 equivalents per amine) for 5 hours at RT under constant agitation (800 rpm). The HC protein (1 mg/mL in 0.1 M phosphate buffer, pH 8) was then added to the dendriplex suspension, and the mixture reacted overnight with continuous agitation (800 rpm) at RT (HC final concentration of 0.25 mg/mL, pH 7.4). After the overnight incubation, the solution, containing the Tg dendriplexes, was purified using an Amicon Ultra-0.5 Centrifugal Filter Unit with a 50 kDa cut-off (Millipore).

### 2.4. Physicochemical characterization of non-targeted (nTg) and targeted (Tg) dendriplexes

Tg or non-targeted (nTg) siPTEN-dendriplexes (N/P 5, 10 or 20) were prepared as described in Section 2.3. The siPTEN-dendriplexes were then characterized regarding their efficiency of siRNA complexation (SYBR® Gold assay), size and polydispersity index (dynamic light scattering), surface charge (electrophoretic scattering), morphology (transmission electron microscopy, TEM) and the concentration of NP in solution by NP tracking analysis (NTA). Furthermore, the siRNA release capacity was evaluated through the heparin assay.

#### 2.4.1. Complexation efficiency of siRNA

The assessment of siPTEN complexation capacity was conducted using the SYBR® Gold assay as previously described [23]. The results are presented as the relative percentage of complexation, wherein 100% indicates complete siRNA complexation, and 0% represents the absence of any nucleic acid binding. To remove any potential background fluorescence, dendritic copolymer, and single HC protein were used as controls.

#### 2.4.2. Heparin assay

To explore the release of siRNA induced by heparin, nTg and Tg dendriplexes were prepared at N/P ratio 5, 10 or 20 and subsequently incubated with heparin (Sigma-Aldrich) at different concentrations ranging from 0 to 0.5 mg/mL. The incubation was carried out for 2 hours at 37 °C [19]. Following this interaction, the SYBR® Gold intercalation assay was performed as indicated in 2.4.1.

#### 2.4.3. Hydrodynamic size and zeta potential measurements

The hydrodynamic size and polydispersity index (PDI) of assembled Tg dendriplexes were determined as previously described using a Malvern Zetasizer Nano ZS (Malvern Instruments, UK) [23]. The Zetasizer Software (version 7.13) was used for analysis.

#### 2.4.4. Nanoparticle tracking analysis (NTA)

N/P 5 and 10 Tg siPTEN-dendriplexes were analysed as previously described by us [28]. Raw data of laser scattering and particle movement were analysed using the NTA software (version 3.4; NanoSight Ltd, UK). Automatic settings were selected for minimum expected particle size, blur, and minimum track length. The detection threshold was set to 4 and sample viscosity was set to the corresponding viscosity for water (at 25 °C) and temperature to 25 °C. The output data was presented as particle concentration (number of particles per mL).

#### 2.4.5. Transmission electron microscopy (TEM)

N/P 10 nTg and Tg siRNA-dendriplexes were imaged as previously described by us using a Jeol JEM 1400 operated at 80 kV [23]. Images were processed using ImageJ software (version 1.54d; National Institutes of Health, USA).

### 2.5. Cell culture

The culture of mouse neuroblastoma and rat neuron hybrid (ND7/23, ECACC), mouse hippocampal neuronal (HT22, provided by Dave Schubert at the Salk Institute), and mouse embryonic fibroblast (NIH 3T3, ECACC) cell lines was carried out using Dulbecco’s Modified Eagle’s Medium (DMEM) with GlutaMAX™ (Gibco). The culture medium was supplemented with 10% (v/v) heat-inactivated (56 °C, 30 minutes) foetal bovine serum (FBS) and 1% (v/v) antibiotic solution containing 10 000 U/mL penicillin and streptomycin (P/S), all provided by Gibco. The cells were maintained at 37 °C in a 5% CO_2_ environment. To ensure the absence of mycoplasma contamination, cell lines were regularly examined using polymerase chain reaction (PCR) analysis. The cells used for the experiments were between passages 1 and 15 after thawing from liquid nitrogen.

Motor neuron cultures were prepared as described by Ionescu et al. (2022) [33]. Cells were then seeded and incubated in the medium previously detailed in this Section.

Primary mouse neuronal cortical cells were isolated from the prefrontal cortex of E16.5 wild-type mouse embryos. The cerebral cortices were carefully dissected and incubated with trypsin (1.5 mg/mL, Gibco) in Hanks’ Balanced Salt Solution (HBSS pH 7.4, Sigma) without CaCl_2_ and MgSO_4_ at 37 °C for 12 minutes. Subsequently, the medium was removed, and 10% (v/v) FBS-containing HBSS was added. The resulting cell clusters were gently washed three times with non-supplemented HBSS, followed by the addition of Neurobasal Medium (Gibco) supplemented with B27 1× (Gibco), 2% (v/v) horse serum (Gibco), 1% (v/v) P/S, 0.025 mM GlutaMAX™, 25 µM beta-mercaptoethanol (Sigma), 1 µM glial cell line-derived neurotrophic factor (GDNF), 0.5 µM ciliary neurotrophic factor (CNTF) and 1 µM brain-derived neurotrophic factor (BDNF) (all factors from Alomone labs). Cells were mechanically dissociated, and the suspension was filtered using a 70 µm nylon cell strainer (Corning Falcon™). The cells were then seeded (cell density specified in the respective subsection) and finally incubated at 37 °C, 5% CO_2_. The purity of the cortical neurons culture was determined to be above 97% through image analysis using Ilastik and Cell Profiler software (data not shown), after immunocytochemistry with the following antibodies: rabbit anti-glial fibrillary acid protein (GFAP, 1: 1000 dilution, Abcam), rat myelin basic protein (MBP, 1: 100 dilution, AbD Serotec), rabbit anti-oligodendrocyte transcription factor 2 (Olig2, 1: 100 dilution, Abcam), rabbit anti-ionized calcium-binding adapter molecule 1 (IBA1, 1: 500 dilution, Wako Chemicals).

### 2.6. Neurospecificity, internalization and transfection assessment

#### 2.6.1. Cellular binding and neurospecificity in cell lines

The neurospecificity of the newly developed Tg dendriplexes was evaluated using flow cytometry. Due to the similar and appropriate physicochemical characteristics, we focused on exploring NPs prepared at N/P ratios of 5. This approach minimizes the addition of material to the cell cultures.

Briefly, ND7/23, HT22 and NIH 3T3 cells were seeded in 24-well tissue culture plates at 2.5 × 10^4^ viable cells (determined by trypan blue assay) per cm^2^ and incubated in supplemented DMEM with GlutaMAX™ for 24 hours until the culture reached 70–80% confluence. Before incubation with NPs, the medium was replaced by non-supplemented DMEM with GlutaMAX™. Tg and nTg dendriplexes prepared at an N/P ratio 5, complexing Cy5-siRNAmi, were added at a final siRNAmi concentration of 100 nM. After 0.5 and 24 hours of incubation, cells were trypsinized, rinsed with PBS 1×, centrifuged and then resuspended in PBS 1× with 2% (v/v) FBS. Subsequently, cells were analysed using a BD Accuri™ C6 flow cytometer (BD Biosciences). The positive cells were quantified by measuring the Cy5 fluorescence signal (more than 10 000 events were collected per sample, with a minimum of 10 000 viable cells). Untreated cells were used as control. FlowJo software (version 10, FLOWJO, LLC) was used to analyse the resulting data.

#### 2.6.2. Downregulation of GFP expression

Primary mouse neuronal motor cells were isolated from the ventral horns of the spinal cord of E12.5 HB9::GFP mouse embryos as previously reported [33]. Briefly, the spinal cords were carefully isolated and incubated with trypsin (25 mg/mL) in PBS 1× at 37 °C for 10 minutes. Then, spinal cords were subjected to 2-3 incubation periods with DNase (1 mg/mL, Sigma) and the fragments were mechanically digested. To purify cells, a BSA cushion (4% w/v BSA solution in Leibowitz medium, Gibco) and OptiPrep density gradient medium (Sigma) were used.

HB9::GFP motor neurons were seeded in 150 µg/mL poly(L-ornithine) and 3 µg/mL laminin-coated 24-well plates at a density of 2.5 × 10^4^ viable cells per cm^2^ (trypan blue assay) and incubated in the supplemented medium detailed in Section 2.5. Dendriplexes with siGFP at N/P 5 were prepared, as described in 2.3. Transfection was performed using 50 μL dendriplexes in 300 μL of culture medium (siGFP final concentration of 100 nM). Non-treated cells and cells treated with Lipofectamine 2000 (L2k, final concentration of 2 µg/mL, Invitrogen) based lipoplexes carrying siPTEN were used as controls. Twenty-four hours later, the medium was substituted by fresh supplemented medium, and cells were incubated for a further 72 hours. Then, cells were rinsed with PBS 1× and subsequently fixed for 15 minutes with PFA (4% (w/v) in PBS 1×). Following cell fixation, cells were incubated in permeabilization buffer (Triton™ X-100, 0.1% (v/v) in PBS 1× pH 7.4) for 30 minutes and then incubated in a blocking buffer solution composed of 1% (w/v) BSA, 0.25% (w/v) goat serum (Sigma Aldrich) in PBS 1× for 1 hour at RT. Cell nuclei were stained with a solution of Hoechst 33 342 in PBS 1× for 10 minutes. VectaShield (Vector Laboratories) was then used to mount coverslips and image acquisition was conducted using an Olympus FV3000 confocal microscope equipped with 20×/0.8, and 60×/1.42 NA objectives. Confocal images of 512 × 512 pixels (zoom 1, 0.622µm per pixel) were analysed using ImageJ (version 1.54d).

### 2.8. Design, fabrication, and characterization of two- and three-compartment microfluidic platforms

We designed a two-compartment microfluidic system to assess the performance of our neuron-Tg fbB dendriplexes in monocultures, namely, to understand the NP intracellular pathway and compatibility with PNS and CNS neurons (Figure 1A). To explore the effect in the axons of CNS neurons, a three-compartment microfluidic platform was also designed (Figure 1B). Furthermore, a PNS-CNS-on-Chip was developed to assess the performance of our NPs in a co-culture of CNS and PNS neurons (Figure 1C and D). The design, fabrication and characterization of the devices are detailed in Support Information (SI).

**Figure 1.**
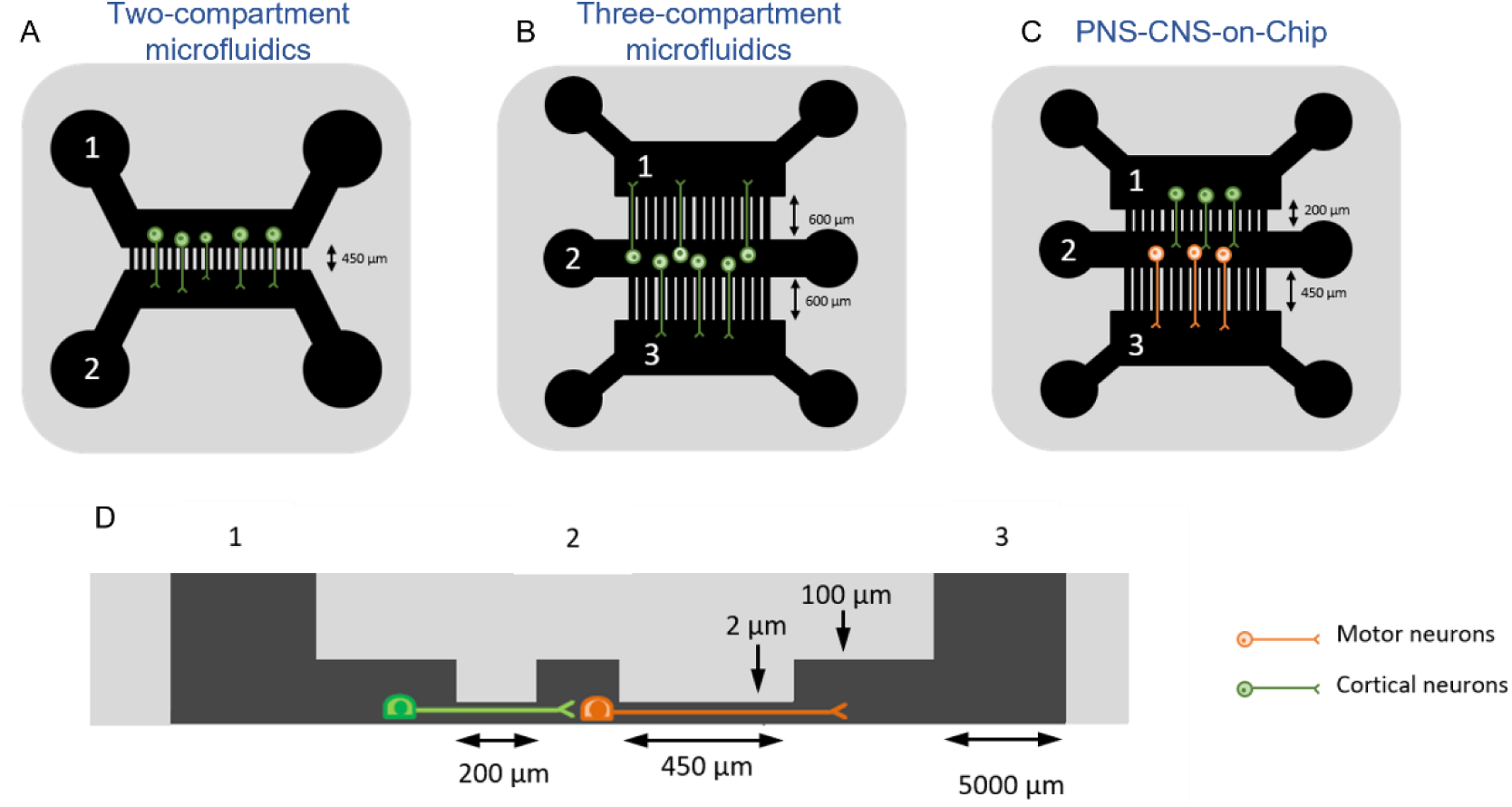
Schematic representation of the microfluidic chips design employed in this study. (A) Two-compartment microfluidics with 450 µm microchannels; (B) Three-compartment microfluidics with two sets of 600 µm microchannels; (C) PNS-CNS-on-Chip, a three-compartment with two distinct sets of microchannels (200 and 450 µm); (D) lateral view of the PNS-CNS-on-Chip with representation of a co-culture of neurons.

#### 2.8.1. Microfluidic design and fabrication

The two and three-chamber microfluidic chips and the PNS-CNS-on-Chip were designed using CleWin4 from WieWeb Software (Hengelo, The Netherlands) and fabricated in poly(dimethylsiloxane) (PDMS) by soft lithography and replica moulding. The cell culture chambers were designed to interconnect through a network of 139 microchannels, each with a cross-Section of 2 × 10 μm and lengths of 200, 450, or 600 µm, allowing axon extension (Figure 1 and Table S2, SI). The 2-μm-high microchannels impede cell migration while allowing axon growth, and the lengths spanning 200 µm or longer ensure complete separation of neuron cell bodies from axonal terminals.

Master moulds were fabricated in silicon wafers with multi-height structures combining deep reactive ion etching (DRIE) and SU-8 photolithography. First, the 2 μm-high microchannels were patterned by direct write laser (DWL) on AZP4100 photoresist and etched directly on the Si substrate by DRIE. To fabricate the larger chambers, SU-8 2100 (MicroChem SU-8 2000 Series) was spin-coated, patterned through a chrome soda lime glass photomask using UV and developed in PGMEA (further details in SI).

SYLGARD™ 184 PDMS Elastomer (10:1 mix of silicone elastomer and its curing agent, Dow Corning) was degassed and cured over the positive relief master for 2 hours at 70 °C. Inlets and outlets acting as media reservoirs were opened using a steel biopsy punch (Ø 5 mm, KAI medical, BP-50F). Following sterilization through a wash with 70% (v/v) ethanol and 10 minutes of exposure to UV, the PDMS stamps were air plasma treated (Tergeo plasma cleaner TG100, Pie Scientific) and bonded to glass coverslips (22 × 22 mm, Normax). The glass coverslips were previously coated with poly(D-lysine) (PDL, 50 µg/mL in 0.1 M borate buffer pH 8.5, Sigma-Aldrich) for primary cortical neurons or poly-L-ornithine (PLO, 1.5 µg/mL in PBS 1× pH 7.4, Sigma-Aldrich) and laminin (5 μg/mL in autoclaved water, L2020, Sigma) for primary motor neurons.

#### 2.8.2. Microfluidic characterization and simulation of fluid movement

The PDMS microfluidic chips were fully characterized through scanning electron microscopy (SEM) and AFM. The channel and microchannel topography was inspected using a Quanta 400 FEG microscope (FEI Company, USA) following a coating with a gold-palladium layer (SPI Supplies). Additionally, AFM was done with a PicoPlus scanning probe microscope, interfaced with a Picoscan 5000 controller (Keysight Technologies, USA) and PicoView 1.20 software (Keysight Technologies, USA), coupled to an inverted optical microscope (Observer Z1, Zeiss, Germany). All measurements were performed at RT using a triangular tip (MLCT-BIO-DC probe, Bruker) in contact mode.

##### Fluid Flow Simulation and Validation

The fluid flow between two large culture chambers was estimated. First, the hydraulic resistance of each channel was calculated using equation (1):

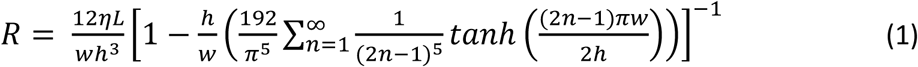

where R is the hydraulic resistance (Pa·s m^-3^), η the fluid viscosity (N·s m^-2^), and L, w and h are the channel dimensions (m). The total resistance of the 139 parallel microchannels was calculated by summing parallel resistors. Furthermore, to estimate the flow rate, the Hagen-Poiseulle law was used (equation 2):

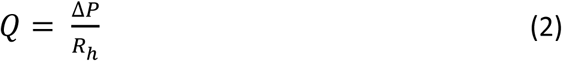

where Q is the flow rate (m^3^·s^-1^), ΔP is the pressure difference across the two main culture chambers (Pa) and R_h_ is the hydraulic resistance of the channels. In this work, the pressure difference is maintained hydrostatically by the addition of different volumes of culture media to each chamber and can be calculated using (equation 3):

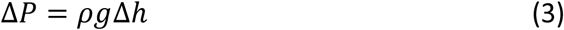

where ρ is the fluid density, g is the gravitational acceleration and Δh is the height difference between the two columns of liquid at either side of the channel. Given the interdependency between the height of the media columns, the pressure difference and the flow rate, their evolution through time was further estimated using a MATLAB script (Mathworks, Inc. Natick, MA, US).

### 2.9. Exploring the intracellular movement of dendriplexes in microfluidic models

#### 2.9.1. Dendriplexes intracellular trafficking

Live retrograde axonal transport of NPs was assessed using the microfluidic-based culture systems. Cortical or motor neurons were seeded in the proximal compartment (cell soma compartment) of a two-compartment microfluidic device (Figure 1A) at a density of 25 × 10^4^ viable cells (trypan blue assay) in 0.161 cm^2^ (total area of the culture chambers). On the 5^th^ day post-seeding *in vitro* (DIV5), Tg and nTg N/P 5 dendriplexes were prepared with Cy5-labeled siRNAmi and added to the axonal compartment, at a final siRNAmi concentration of 100 nM. NPs only contact the neuron axon terminals (Figure 1A, compartment 2) to evaluate their capacity to be internalized and reach the cell body. Untreated cells were used as controls. To establish a hydrostatic pressure difference and create a slow but continuous counter-flow through the microchannels, a lower volume of media (50-μL difference) was added to the axonal compartment, thus guaranteeing unilateral flow between the cell soma and axonal compartments. After a 16-hour incubation period, axons were imaged live using a Crest X-Light V3 spinning disk confocal microscope (Nikon) in a 37 °C heated chamber. In each treatment group, a minimum of 24 microchannels, each containing at least one axon, were randomly selected for live cell imaging. Half an hour before the acquisition, cells were incubated with CellMask™ Plasma Membrane Stain 488 (1: 1000 dilution, C37608, Invitrogen). Images were captured for at least 5 minutes at a frame rate of 0.33 FPS at a resolution of 1024 × 1024 pixels using a Plan Apo VC 60xA WI DIC N2 water objective (numerical aperture: 1.2). Images were acquired from three independent experiments. Subsequent image analysis was conducted using Fiji ImageJ 1.54f [33].

#### 2.9.2. Intracellular dynamics assessment

Cell seeding and treatment with dendriplexes followed the procedure outlined in the previous subsection. At specific time points (8, 12, 24, 48, and 120 hours), cells were fixed with microtubule-protecting (MP) fixative buffer. MP-PFA solution consists of 4% (w/v) PFA, 65 mM 1,4-Piperazinediethanesulfonic acid (PIPES), 25 mM 4-(2-hydroxyethyl)-1-piperazineethanesulfonic acid (HEPES), 10 mM ethylene glycol-bis(β-aminoethyl ether)-N,N,N′,N′-tetraacetic acid (EGTA), and 3 mM MgCl_2_ in PBS 1× (all acquired from Sigma-Aldrich). The fixation process comprised three steps: Initially, MP-PFA was added to the cell medium in the reservoirs (1:1 ratio) and incubated for 15 minutes. Then, the mixture was removed, and MP-PFA was added to fully cover the reservoirs. Finally, MP-PFA was aspirated from the reservoirs, the microfluidic device was detached from the coverslip, and MP-PFA was directly applied to the coverslip, which was incubated for an additional 15 minutes. Following cell fixation, cells were permeabilized (Triton™ X-100, 0.1% (v/v) in PBS 1×) for 15 minutes and then incubated in a blocking buffer composed of 1% (w/v) bovine serum albumin (BSA, NZYTech) in PBS 1× for 45 minutes at RT. Subsequently, cells were rinsed with PBS 1×, and then incubated with mouse anti-βIII tubulin (2 µg/mL) overnight at 4 °C. After incubation with the primary antibody, samples were incubated with Alexa Fluor 488 donkey anti-mouse IgG (H+L) secondary antibody (2 µg/mL) for 1 hour at RT and then washed again with PBS 1×. Finally, cell nuclei were stained with a solution of Hoechst 33 342 (5 µg/mL, Thermo Fisher Scientific) in PBS 1× for 10 minutes. Fluoromount™ Aqueous Mounting Medium (Sigma) was then applied to mount the coverslips on the microscope slides. Image acquisition was performed using a Leica TCS SP5 Confocal Microscope (Leica Microsystems, Inc.), using an HCX PL APO CS 63.0×1.40 oil objective. Confocal image z-stacks of 1024 × 1024 pixels (16-bit depth and zoom levels between 1.7 and 2.5, voxel size: 240.5 nm) were captured. For Hoechst, Alexa Fluor 488 and Cy5 acquisition, the following SP mirror channels were set to 414-486 nm, 497-589 nm, and 695-765 nm, respectively. Images were then processed using the Leica Application Suite X software (Leica Microsystems).

### 2.10. Biocompatibility profile of dendriplexes assessed in microfluidic neuronal cultures

The impact of the fbB dendrimer and its corresponding Tg and nTg dendriplexes on embryonic mice motor neurons and cortical neurons was assessed using the lactate dehydrogenase (LDH) method and microelectrode arrays (MEAs). With both methods, we explored the effect of these dendritic NPs on well plates and in microfluidic cultures.

#### 2.10.1. LDH cytotoxicity assay

After seeding cells in two-compartment microfluidics as described in Section 2.9.1, cells were exposed to the fbB dendrimer, Tg or nTg dendriplexes, (added only on the axonal compartment) for 24 hours. Subsequently, the cell culture medium was collected from the 5 mm reservoirs from the cell soma compartment, and the released LDH was quantified using the CyQUANT™ LDH cytotoxicity assay kit (Thermo Fisher) following the manufacturer’s guidelines. The absorbance of the final product (red formazan) was measured at λ = 490 nm using a BioTeK® Synergy MC multi-plate reader (Winooski, VT, USA). The same study was also conducted using a 24-well plate, as previously described by us [28], to enable comparative analysis of the observed effects. As controls, cells were also treated with L2k (final concentration of 2 µg/mL)-siPTEN lipoplexes, or Triton™ X-100 0.1% (v/v) in PBS 1× pH 7.4. The data are presented as the percentage of LDH released by treated cells relative to non-exposed cells.

#### 2.10.2. Neuronal electrical activity

##### Assessment in well-plate MEAs

Commercial six-well-plate MEAs (60-6wellMEA200/30iR-Ti, Multichannel Systems), each containing a recording area of 3 × 3 grids of embedded gold electrodes (30-µm diameter, 200-µm pitch), were used to monitor the electrical activity of neuronal networks non-invasively, at different time points. The day before cell seeding, MEAs were coated as described in Section 2.7. Then, 20 × 10^4^ viable (trypan blue assay) primary cortical or motor neurons were seeded in each well. Cells were grown in an incubator at 37°C and 5% CO_2,_ and 50% of the medium was refreshed every 2 days. At DIV5, dendriplexes were prepared and added to cells as described in Section 2.7. At DIV12, spontaneous neuronal activity (10 kHz sampling rate, 5-minute recording), was recorded upon mounting the MEA plate into the recording system. Detected spikes were sorted and analysed using a MATLAB script (The Math-Works, Inc., Natick, MA). Untreated cells and cells treated with Triton X-100 0.1% (v/v) in PBS 1× pH 7.4 were used as control.

##### Assessment in microfluidic platforms integrating MEAs

Commercial planar MEAs (60MEA200/30iR-ITO, Multichannel Systems) comprising sixty electrodes (30-µm diameter, 200-µm pitch) were used to monitor the electrical activity of neuronal networks in the PNS-CNS-on-Chip (Figure 1C). Prior to usage, both the microfluidics and MEAs were briefly submerged in 70% ethanol, followed by air-drying inside a laminar flow, and UV sterilization. Subsequently, microfluidic devices were carefully assembled on top of the MEAs under the guidance of a stereomicroscope, ensuring precise alignment of the microchannels with the microelectrodes. The platform was then coated as described in Section 2.7. In this assay, a three-compartment platform to create two distinct cultures of different neurons (PNS and CNS neurons) was used. The cell bodies and axon terminals were physically separated by sets of microchannels, measuring 200 and 450 µm (Figure 1C). In the three-compartment design, primary motor neurons were seeded in the central compartment (Figure 1C, compartment 2) at 20 × 10^4^ viable cells (trypan blue assay) in 0.076 cm^2^ (total area of the culture chamber) and incubated for 24 hours. Then, primary cortical neurons (25 x 10^4^ viable cells, (trypan blue assay) in 0.076 cm^2^) were seeded in the lateral compartment (Figure 1C, compartment 1), being the two different cultures of neurons separated by 200-µm microchannels (Figure 1D). Throughout the experiment, a hydrostatic pressure difference was maintained to ensure appropriate axonal growth direction. At DIV5, dendriplexes were prepared as described in Section 2.3 and added to cells in the free compartment (Figure 1C, compartment 3). At DIV12, spontaneous neuronal activity (10 kHz sampling rate; 5-minute recording), was recorded upon mounting the MEA plate into the recording system. Non-treated cells and cells treated with Triton™ 0.1% (v/v) in PBS 1× pH 7.4 were used as controls. Detected spikes were sorted and analysed using a MATLAB script and SpikeSorter software (version 5.20, Vancouver, BC Canada) [34].

### 2.11. Axonal outgrowth

To assess the biological effect through axonal outgrowth analysis, three-compartment microfluidic devices were used (Figure 1B). Primary cortical neurons were seeded in the middle compartment (microchannels: 600/600 µm, Figure 1B, compartment 2) at a density of 20 × 10^4^ viable cells (trypan blue assay) in 0.161 cm^2^. On the DIV5, Tg and nTg N/P 5 dendriplexes were prepared with siPTEN and added to the cell soma compartment (Figure 1B, compartment 2), at a final siRNA concentration of 100 nM. Cells treated with free dendrimer only, siPTEN, and untreated cells were used as controls. At DIV10, cells were fixed with MP-PFA following the 3-step fixation procedure described in Section 2.9.2. Following cell fixation, cells were incubated in permeabilization buffer (Triton™ X-100, 0.1% (v/v) in PBS 1×) for 15 minutes and then incubated in a blocking buffer composed of 1% BSA in PBS 1× for 1 hour at RT. Subsequently, cells were rinsed with PBS 1×, and then incubated with rabbit anti-Neurofilament H antibody (0.6 µg/mL, ab207176, Abcam) overnight at 4 °C. After primary antibody incubation, samples were incubated with Alexa Fluor 488 donkey anti-rabbit IgG (H+L) secondary antibody (2 µg/mL, A-21206, Invitrogen) for 1 hour at RT and then washed again with PBS 1×. Finally, cell nuclei were stained with a solution of Hoechst 33 342 (5 µg/mL, Thermo Fisher Scientific) in PBS 1× for 10 minutes. Untreated cells were used as controls. Fluoromount™ Aqueous Mounting Medium (Sigma) was then applied to mount the coverslips. The axonal outgrowth within the axonal compartments (Figure 1B, compartments 1 and 3) was assessed using linear Sholl analysis MATLAB scripts, kindly provided by Martin Offterdinger and Prof. Rüdiger Schweigreiter [35].

### 2.11. Inter-neuronal migration capacity evaluation

Primary motor neurons were seeded in the middle compartment PNS-CNS-on-Chip (microchannels: 450/200 µm, Figure 1C, compartment 2) at a density of 25 × 10^4^ viable cells (trypan blue assay) in 0.161 cm^2^. Twenty-four hours later, primary cortical neurons were seeded in the lateral compartment connected by 200 µm microchannels (Figure 1C, compartment 1) at a density of 25 × 10^4^ viable cells (trypan blue assay) in 0.161 cm^2^. On the DIV5, Tg and nTg N/P 5 dendriplexes were prepared with Cy5-siRNAmi and added to the free compartment, at a final siRNA concentration of 100 nM. Untreated cells were used as control. After 120-hour incubation, cells were fixed with MP-PFA (3-step fixation procedure described in Section 2.9.2). Following cell fixation, cells were incubated in permeabilization buffer (Triton™ X-100, 0.1% (v/v) in PBS 1×) for 15 minutes and then incubated in a blocking buffer composed of 1% BSA in PBS 1× for 1 hour at RT. Subsequently, cells were rinsed with PBS 1×, and then incubated with mouse anti-βIII tubulin (2 µg/mL) overnight at 4 °C. After incubation with the primary antibody, samples were incubated with Alexa Fluor 488 donkey anti-mouse IgG (H+L) secondary antibody (2 µg/mL) for 1 hour at RT and then washed again with PBS 1×. Finally, cell nuclei were stained with a solution of Hoechst 33 342 (5 µg/mL, Thermo Fisher Scientific) in PBS 1× for 10 minutes. Fluoromount™ Aqueous Mounting Medium (Sigma) was then applied to mount coverslips. Image acquisition was performed using a Leica TCS SP5 Confocal Microscope (Leica Microsystems, Inc.), using HCX PL APO CS 63.0×1.40 oil objective. Confocal image z-stacks of 1024 × 1024 pixels (16-bit depth and zoom levels between 1.7 and 2.5, voxel size: 240.5 nm) were captured. For Hoechst, Alexa Fluor 488 and Cy5 acquisition, the following SP mirror channels were set to 414-486 nm, 497-589 nm, and 695-765 nm, respectively. Images were acquired from three independent experiments. Images were then processed using Leica Application Suite X software (Leica Microsystems) and Fiji ImageJ 1.54f.

### 2.12. Statistical Analysis

Data analysis and presentation were performed with IBM SPSS Statistics version 26 (SPSS Inc., USA) and GraphPad Prism version 10 for Windows (GraphPad Software, USA). The normal distribution of data was confirmed using the Shapiro-Wilk test. Differences were evaluated through one-way or two-way ANOVA, followed by Tukey’s multiple comparisons tests, as detailed in the results. Statistical significance was indicated by P values less than 0.05.

## 3. Results and Discussion

### 3.1. Development and physicochemical characterization of targeted dendriplexes for neuronal siRNA delivery

To attribute neurotropism to our fbB dendriplexes, these were functionalized with the non-toxic and neurospecific TeNT HC domain. Our Tg dendriplexes were engineered to mimic the TeNT cell entry mechanisms and gain access to the peripheral neuron cell bodies through active retrograde axonal transport and from there reach CNS neurons [31, 36].

To obtain the Tg NPs, firstly nTg dendriplexes formed between fbB dendrimers and a therapeutic siRNA (siPTEN) were prepared at N/P 5, 10 or 20, following the procedure previously described by us (Figure 2) [19, 28]. Afterward, the dendriplexes were functionalized, via their free amines, with a 1 kDa heterofunctional linker (MAL-PEG-NHS) to subsequently enable binding of the HC domain via thiol groups. This functionalization with HC was carried out by incubation of the dendriplexes with the TeNT fragment. A thiol-ene Michael addition between the thiol at the HC cysteines and the MAL present in the mentioned linker on the surface of dendriplexes led to the Tg dendriplexes (Figure 2C) [37].

**Figure 2.**
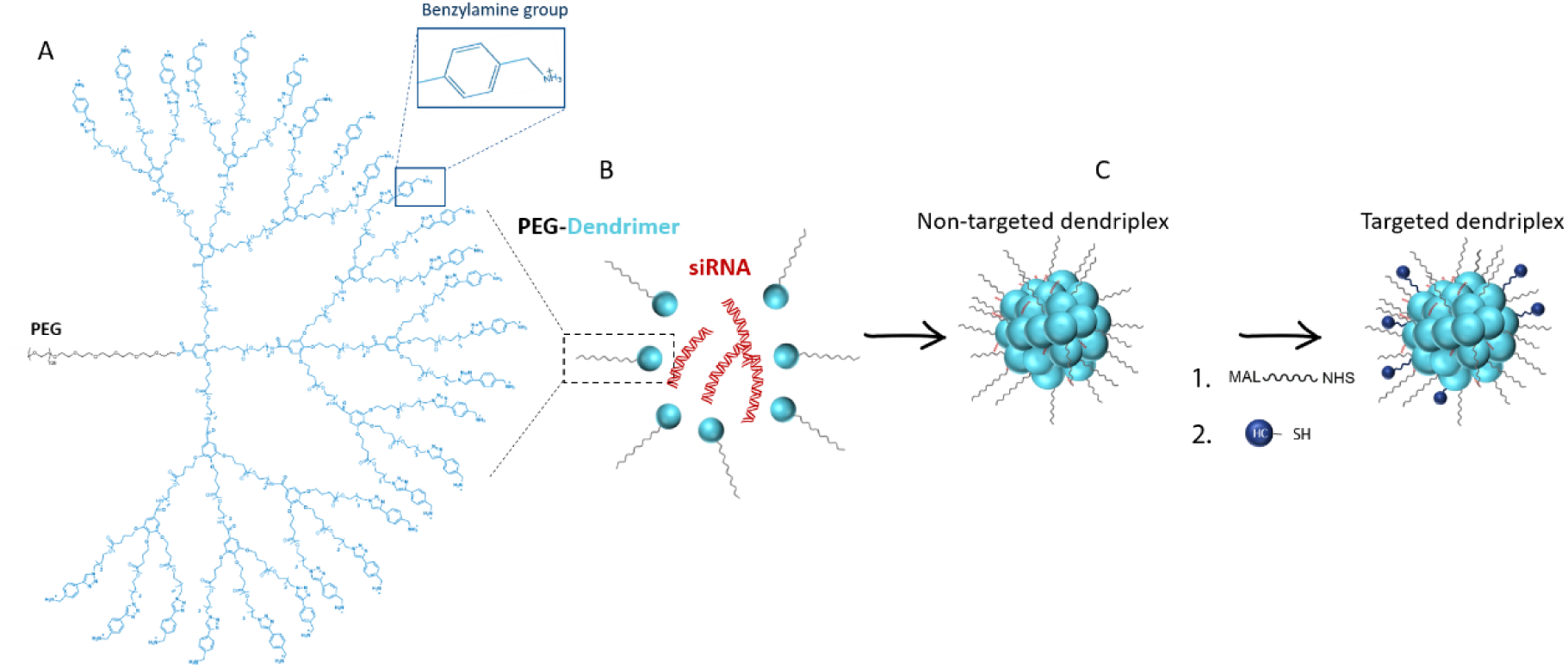
Preparation of non-targeted (nTg) and targeted (Tg) siRNAdendriplexes. (A) Structure of the generation 3 (G3) of fully biodegradable benzylamine-terminated PEG–GATGE (poly(Ethylene Glycol)-Gallic Acid-Triethylene Glycol Ester) dendritic block copolymers; (B) Schematic representation of the preparation of nTg siRNA dendriplexes and (C) Tg dendriplexes, functionalized with HC.

The complexation capacity of the positively charged vector with the negatively charged siPTEN as a function of the N/P ratios was assessed by exploring a nucleic acid dye (SYBR® Gold) accessibility assay for both Tg and nTg NPs (Figure 3A). The percentage of complexed nucleic acid significantly increased with the N/P ratio. Remarkably, for all N/P ratios tested, the fbB dendrimers’ ability to complex siRNA was above 68% and 73% for the nTg and Tg N/P 5 dendriplexes, respectively, reaching values of 88-89 % for the highest evaluated N/P ratios. The complexation efficiency of both Tg and nTg very similar, suggesting that the surface functionalization with the TeNT protein fragment does not affect the complexation efficiency of our fully biodegradable vector.

**Figure 3.**
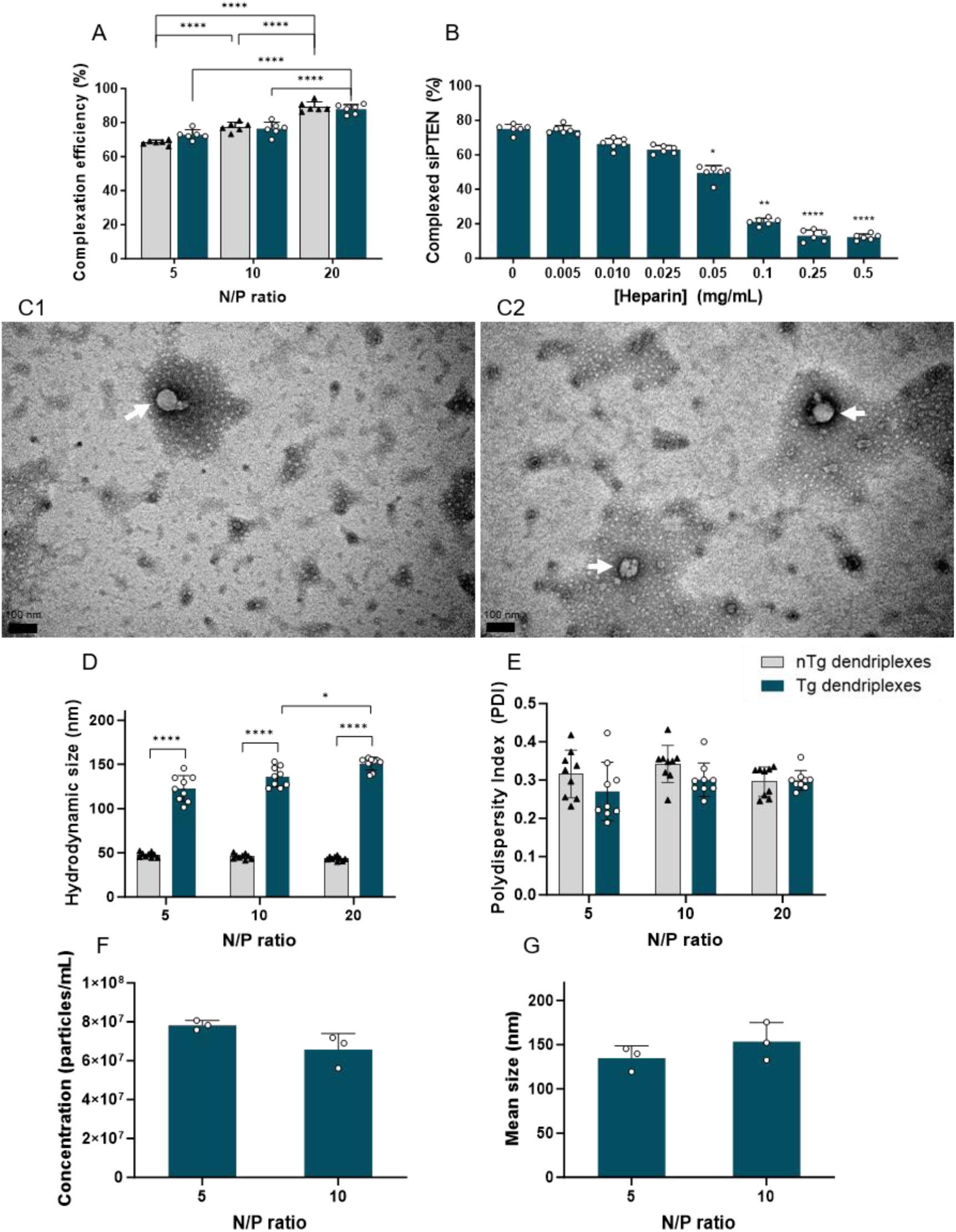
Physicochemical characterization of targeted (Tg) and non-targeted (nTg) dendriplexes prepared with siPTEN as a function of N/P ratio. (A) SYBR® Gold exclusion assay (room temperature); (B) Heparin dissociation assay of N/P 10 Tg dendriplexes; (C) Transmission electron microscopy (TEM) images of (C1) nTg N/P 5 dendriplexes and (C2) Tg N/P 5 dendriplexes. White arrows indicate dendriplexes. Scale bar: 100 nm; (D) Size distribution (Dynamic Light Scattering, DLS); (E) Polydispersity index (PDI) (DLS); (F) Concentration of particles in solution (NP tracking analysis, NTA); (G) Mean size of particles in solution (NTA). Results are shown.

The assessment of an effective delivery vector goes beyond just its complexation capacity. A vector’s proficiency in releasing its payload within cells is equally crucial. Therefore, to gauge the assembly reversibility of Tg and nTg dendriplexes (Figure S6, SI and 3B, respectively), the prepared complexes were incubated with heparin at 37°C. At physiological pH, heparin is an anionic polymer widely used for assessing the destabilization and release of nucleic acids from delivery nanosystems [38]. The nTg dendriplexes were prepared at both the highest and lowest ratios (5 and 20), while Tg dendriplexes were prepared at N/P 10. All of them were subjected to the heparin disassembly test, exhibiting a notable release of siPTEN. Particularly noteworthy was the statistically significant release at heparin concentrations above 0.05 mg/mL, with approximately 50% of the siPTEN released. Remarkably, at the two highest concentrations (0.25 and 0.5 mg/mL), 90% of the siPTEN was released. Our N/P 5 and N/P 20 nTg dendriplexes also showcased their ability to release the siPTEN (Figure S6, SI), evidenced by their strong siRNA release when competing with heparin (at concentrations up to 0.5 mg/mL). This demonstrates that the targeting functionalization with the TeNT protein fragment does not alter the siRNA release capacity, specifically of the siPTEN (Figure S6, SI).

The morphology, hydrodynamic size, polydispersity index (PDI), and concentration of the nTg and Tg dendriplexes were determined by transmission electron microscopy (TEM), dynamic light scattering (DLS), and NP tracking analysis (NTA), respectively (Figure 3). TEM images showed that the developed nTg and Tg dendriplexes presented similar spherical morphologies (Figure 3C). Both dendriplexes formulations (nTg and Tg) presented small average sizes at the N/P ratios tested (Figure 3D). Noteworthy, although the Tg dendriplexes were significantly larger than their nTg counterparts, these Tg dendritic NPs still maintain significantly low sizes suitable for the intended neuronal application [39]. Moreover, both dendriplex populations displayed a consistent homogeneity in sizes, with PDIs below 0.35 in the case of nTg dendriplexes and below 0.30 for Tg dendriplexes (Figure 3E). Regarding the concentration of NPs in solution for Tg dendriplexes (Figure 3F), a slight decrease in particle concentration was observed with an increasing N/P ratio (from 5 to 10), but not statistically significant. NTA also facilitated the determination of the mean particle size in suspension (Figure 3G). Tg dendriplexes at N/P ratios of 5 and 10 exhibited sizes ranging from 134.9 to 153.5 nm, which closely align with those determined by DLS, confirming the values obtained through this technique.

### 3.2. Biocompatible profile of dendriplexes in conventional neuronal culture

To evaluate neurons’ viability after treatment with the proposed nanosystems, cells were cultured in conventional well plates and incubated for 48 hours with nTg and Tg N/P 5 dendriplexes carrying siPTEN. Then, the LDH cytotoxicity assay was performed (Figure S7, SI). Both Tg and nTg NPs had a minimal impact on the cell membrane and this determination is drawn from the low percentage of released LDH observed in embryonic motor neurons (Figure S7A, SI) and cortical neurons (Figure S7B, SI). The lipid vector used as a control (L2k), considered one of the gold standards for *in vitro* transfection, exhibited a significantly greater impact on the membrane of both types of neurons, approaching the LDH release value achieved by the cytotoxic Triton-X 100.

Furthermore, the impact of siPTEN-loaded nTg and Tg dendriplexes on neurons is innovatively assessed through the analysis of their electrical activity, providing a novel approach to evaluating biocompatibility. This pioneer assay examines the effect of NPs by measuring the electrical activity of neurons, crucial for determining biological applicability (Figure S8, SI). For NPs to be considered biocompatible, they should not disrupt the normal electrophysiological response of neuronal cells, as it could hinder neuronal communication. The impact of our formulations was evaluated in well plates by administering NPs to the entire cell population (Figure S7C and S7D, SI). After measuring and analyzing the electrical activity, nTg and Tg dendriplexes exhibited electrical features (number of spikes, number of bursts, interspike interval (ISI), and mean intra-burst interval (IBI)) similar to untreated cells, signifying no electrophysiological changes in the PNS or CNS neuronal population. Both formulations yielded much more promising results than the lipid vector, which significantly impacted the axonal network of motor and cortical neurons (Figure S7C and D). Additionally, the neuronal spiking data was similar between dendriplexes treated and untreated samples (Figure S8M, SI). Most electrodes were active, showing a heterogeneous firing-rate distribution (Figure S8N and O, SI).

### 3.3. Revealing the neurospecificity and efficient siRNA delivery of targeted dendriplexes

After confirming the biocompatibility of our vectors, the neurotropism of our Tg dendriplexes towards both neuronal cells from the PNS and CNS was assessed.

Initial tests were conducted in cell lines, where ND7/23 and HT22 cells were exposed to either nTg or Tg dendriplexes (N/P ratio of 5) carrying a Cy5-labeled siRNAmi for 0.5 and 24 hours. Cell membrane interaction was determined by flow cytometric analysis (Figure 4A and 4B). NIH 3T3 fibroblast cells were used as a negative control. After half an hour of incubation, Tg dendriplexes exhibited statistically significant higher interaction with neuronal cells (ND7/23 and HT22) compared to the nTg dendriplexes (Figure 4A). Notably, approximately 94% of ND7/23 cells showed a Cy5 fluorescence signal following treatment with the Tg NPs, whereas the signal was observed in 79% of cells treated with the nTg dendriplexes. In HT22 cells, the incubation with the Tg dendriplexes resulted in 95% of cells displaying a fluorescence signal, in contrast to the 75% of cells incubated with the nTg dendriplexes. At this time point, the most notable difference was in the percentage of positive cells, with no significant differences recorded in fluorescence intensity after 0.5 hours of incubation. On the other hand, after 24 hours of incubation (nearly 100 % Cy5-positive cells), variations in cell-associated fluorescence intensity also became apparent (Figure 4B). When ND7/23 and HT22 were exposed to Tg dendriplexes, the cells exhibited significantly higher fluorescence values (8.4 × 10^4^ and 7.2 × 10^4^, respectively) compared to the intensity obtained after incubation with nTg dendriplexes (4.7 × 10^4^ and 2.4 × 10^4^, respectively). The presence of Cy5-siRNAmi within the cells, increased by factors of 1.8 in ND7/23 cells and 3.0 in HT22 cells. The interaction of our Tg NPs was also statistically significantly superior in neuronal cells compared to the non-neuronal cells (NIH 3T3) in both time points tested (0.5 and 24 hours). The nTg dendriplexes interacted with the NIH 3T3 cells at a commendable level, with Cy5 signal detected in 76% of the cells. However, when exposed to the Tg NPs, unlike the interaction increase observed in neuronal cells, approximately 74% of the non-neuronal cells were positive for the fluorescent signal. The similar percentage found with nTg and Tg dendriplexes in NIH 3T3 cells suggests that the functionalization with the neurotargeting moiety might not influence the interaction with non-neuronal cell lines. As anticipated, the functionalization of our dendriplexes with HC contributed to an enhanced binding affinity exclusively with neuronal cells. The higher association of Tg dendriplexes to ND7/23 and HT22 cells compared to NIH 3T3, indicates the efficient targeting ability. Despite leading to higher NP sizes, the functionalization with the TeNT HC domain may have contributed to this enhanced transfection efficiency. Although TeNT is primarily known as a toxin that interacts with the peripheral motor neurons and sensory neurons [36], this study further demonstrates that the binding domain of this toxin also exhibits a significant affinity for CNS neurons. Importantly, the use of this targeting moiety also imparts a tropism for CNS neurons to the NPs.

**Figure 4.**
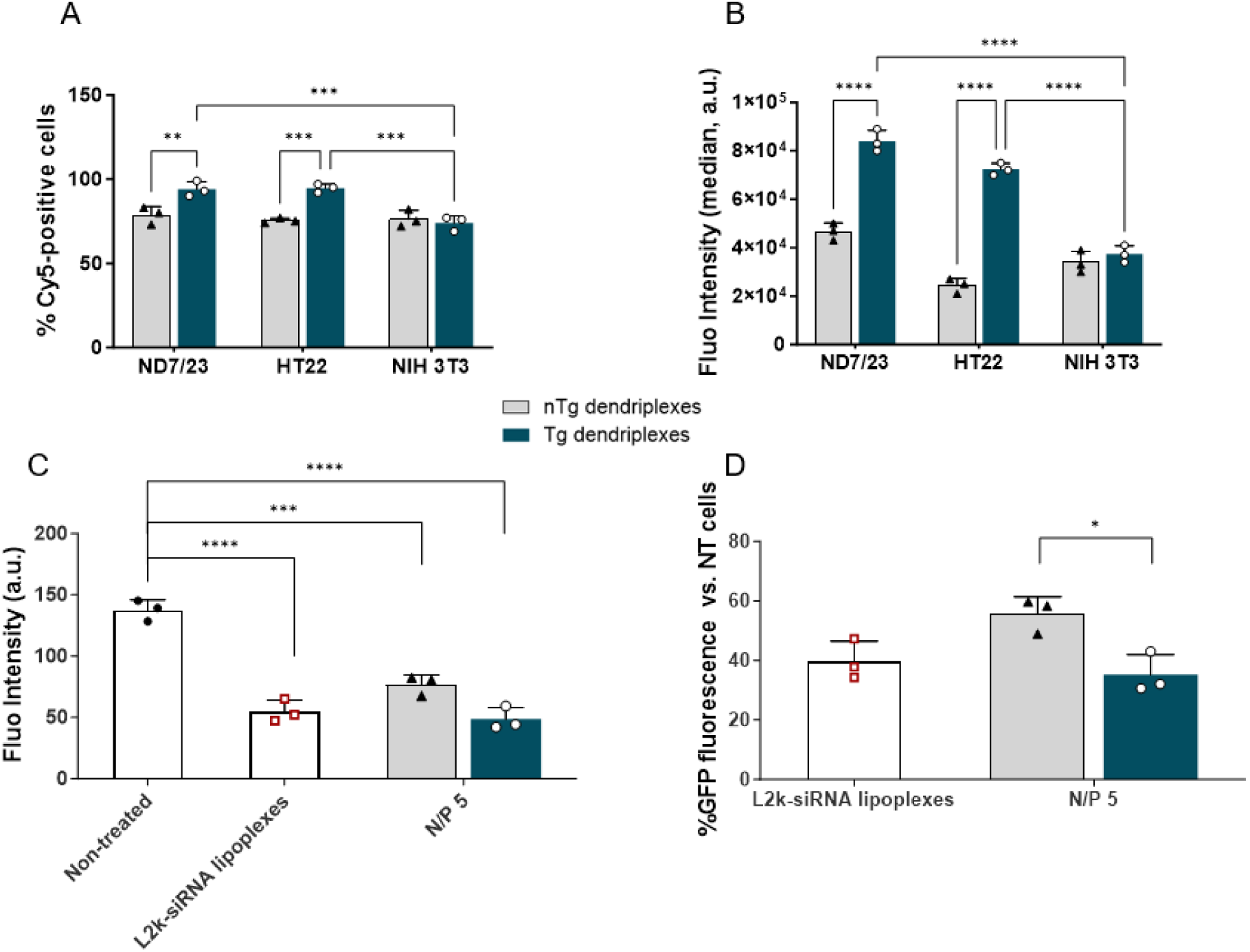
Neurospecifity of targeted dendriplexes. Cellular interaction was assessed by flow cytometry in neuronal (ND7/23 and HT22) and non-neuronal (NIH3T3) cell lines. Cells were incubated with nTg and Tg dendriplexes (N/P 5) carrying Cy5-labeled siRNAmi (final siRNAmi concentration of 100 nM) for (A) 0.5 hours and (B) 24 hours. Characterization of (A) Cy5 positive cells (in %) and (B) Cy5 fluorescence intensity; (C) and (D) downregulation of GFP expression in motor neurons seeded in well-plates, after 96-hour incubation with N/P 5 dendriplexes. Results are represented as mean ± SD of three independent experiments (n = 3), with 2 replicates per experiment. One-way ANOVA tests were used for statistical analysis. Significant differences: *p < 0.05, **p < 0.01, *** p < 0.001 and ****p ≤ 0.0001.

Given the neurospecific association observed in different neuronal cell lines, the capacity of the Tg NPs to mediate the delivery of siRNA leads to a downregulation effect was assessed on a reporter gene - GFP. Neurons expressing the HB9-eGFP fusion protein were incubated for 96 hours with Tg dendriplexes (N/P 5) carrying siGFP (final concentration of 100 nM). Cells treated with nTg particles were also used as controls. The fluorescence intensity of eGFP in primary motor neurons, seeded in well-plates, was then quantified through imaging. The transfection efficiency was determined by the reduction in GFP reporter gene expression. As shown in Figure 4C, both NP formulations caused a reduction in fluorescence intensity in the cells. Notably, Tg dendriplexes achieved a significantly greater reduction, silencing approximately 65% of the expression, in comparison to nTg dendriplexes which led to a 44% reduction in GFP expression (Figure 4D). This demonstrates the suitability of these nanosystems for this application. Additionally, the downregulation effect of Tg dendriplexes was very similar to the outcome observed in cells treated with the commercial vector, which resulted in a 60% reduction in GFP expression. Despite this silencing outcome, the *in vivo* applicability of L2k is limited due to its toxicity (Figure S7 and S8, SI). Therefore, our dendriplexes emerge as a promising option, supported by their demonstrated biocompatibility (Section 3.2) and their great transfection efficiency.

### 3.4. Exploring dendriplexes’ performance in microfluidic-based neuronal cultures

Microfluidics is a powerful technology that here was exploited to enable the cultivation of neuronal cells in separate compartments, thus mimicking their anatomy in the human organism, and facilitating controlled exposure to test compounds [40], This controlled environment is invaluable to assess the efficacy, cytotoxicity, and mechanisms of NP-cell interaction and cellular responses. To explore the performance of our siRNA delivery vector, two- and three-compartment microfluidic devices were used to create controlled environments for neurons, separating the cell soma and axon terminals, thereby mimicking the natural neural architecture (Figure 1 and 5).

**Figure 5.**
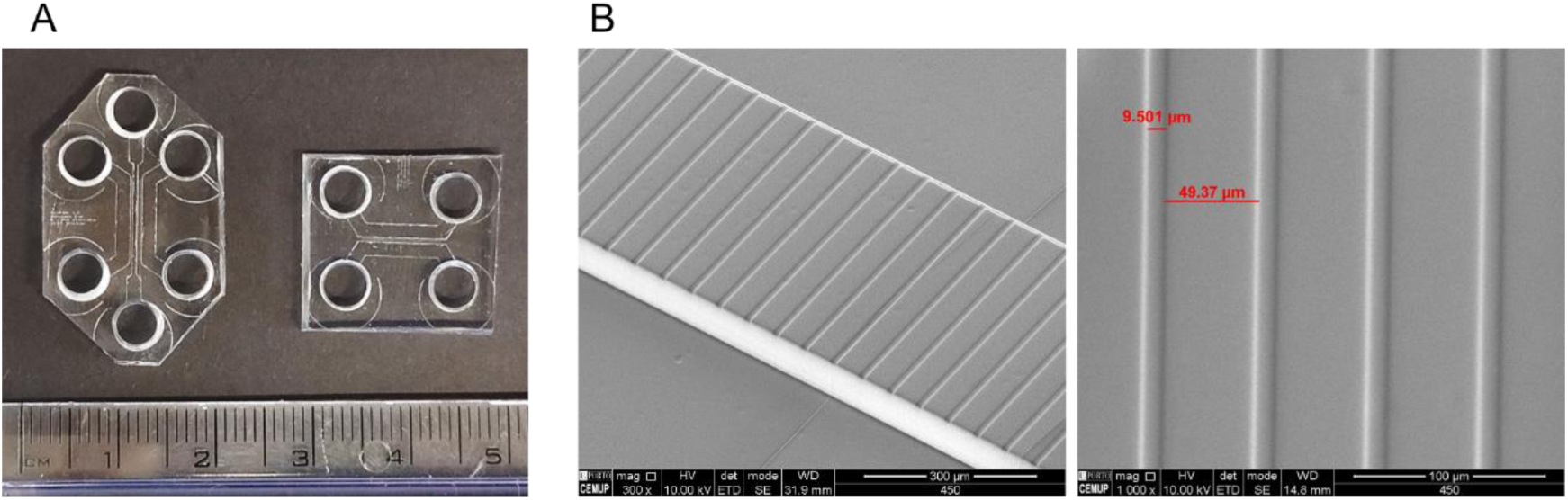
Characterization of two- and three-compartment microfluidic devices. (A) Picture of the poly(dimethylsiloxane) (PDMS) microfluidic devices; (B) Scanning electron microscope (SEM) images of the microchannels.

Our microfluidic platforms (Figure 5A) were produced using PDMS, which is a transparent and biocompatible elastomer material widely used in the field of microfluidics because it is easy to model and can build complex microchannel networks. The produced microfluidic systems were thoroughly characterized by SEM and AFM (Figure 5B, S9 and Table S2, SI).

The two-compartment platform was used for the individual cultures of PNS or CNS neurons (Figure 1A). To create the full PNS-CNS-on-Chip model, we designed a three-compartment platform that enabled the establishment of a co-culture of PNS and CNS neurons in separate culture chambers (Figure 1C). Furthermore, the three-compartment device facilitated the investigation of the axonal growth of CNS neurons (Figure 1B). Altogether, our platforms offered two or three rectangular cell culture chambers, each capable of containing 100–150 μL of cell culture media. The chamber height of 100 µm was crucial to accommodate neurons and provide space for maturation (Figures 1 and 5). One hundred and thirty nice microchannels, each measuring 200, 450, or 600 µm in length and 2 µm in height, connect the culture chambers. Axons are guided and directed through the microchannels, mimicking the natural separation of soma and axons occurring *in vivo*. The 10-µm wide microchannels (Figure 5B) provide controlled communication between compartments, maintaining spatial separation essential for precise control of experimental conditions where axonal terminals and cell bodies can be subjected to selective treatments. Additionally, devices were designed to allow seamless alignment of the microchannel with at least two electrodes from the MEAs to allow the measurement of signal propagation along the axons. To ensure optimal device performance with adequate flow rate and sufficient nutrient supply for cells during culture, the relationship between media volume, height, and flow rate, was estimated using the Hagen-Poiseuille equation and a customized MATLAB script.

To further characterize the devices, the fluid flow was simulated in the compartments and microchannels. To characterize the flow, the hydraulic resistance was first determined (Table S3, SI). The hydraulic resistance of the main- and micro-channels was determined at 1×10^11^ and 7.7×10^16^ Pa·s·m^-3^ respectively. With the total resistance of the 139 microchannels being approximately 5,000 times greater than that of the main culture chamber, even if one were to fill only one of the two chambers at the extremities of the long compartment, just 0.02% of the liquid would flow through the microchannels while the remaining 99.98% would follow the course of the main culture chamber until equilibrium in volumes was reached. All experiments in the two-compartment microfluidics started with 150 μL of cell culture medium in compartment 1 (cell soma compartment) and 100 μL in compartment 2 (axon terminals compartment). It is worth noting that each long compartment has an estimated volume of 1.8 μL and as both long channels are equal, the volume difference between the two remains 50 μL. The 50-μL difference between the two compartments was quickly equilibrated between the two 5 mm inlets/outlets on either side of the cell soma compartment (Table S4, SI). Consequently, each inlet has an excess of 25 μL when compared to the other two chambers on the axon terminals compartment, which corresponds to a height difference of 1.27 mm and a pressure difference of 12.5 Pa. Using Hagen-Poiseuille’s law, a very low flow rate of 22.5 pL/s was estimated through the microchannels. Even if the flow rate were to remain constant throughout the entire duration of the experiment, it would require approximately 13 days, for the volumes in both long channels to equilibrate and for the flow to stop (Figure S10 and S11, SI). This constant counterflow played a pivotal role in our system to maintain chemical gradients, promote axonal growth and guarantee that NP transport was active and not due to diffusion.

#### Axonal retrograde transport of neuron-targeted dendriplexes

Our microfluidic chips were designed as a model to simulate the peripheral administration of neurotropic NPs. This model enables the assessment of the uptake, retrograde axonal transport, and biocompatibility profiles of these NPs in PNS and CNS neurons. As previously commented, to the best of our knowledge, this is the first report employing microfluidic platforms for investigating the pathway and biocompatibility of dendritic NPs in neurons and marks the first report to evaluate the bioperformance of NPs in CNS neurons cultured in microfluidic platforms.

To investigate the potential of dendriplexes to be retrogradely transported from axonal terminals to the neuron cell bodies, live cell imaging experiments were conducted using motor or cortical neuron cultures seeded in the microfluidic devices. Dendriplexes were added to the medium in the axonal terminals compartment, and subsequent NPs transport was imaged in the axons that grew within the microchannels. Live cell imaging analysis revealed the retrograde transport in axons of numerous vesicles loaded with nTg and Tg dendriplexes (N/P 5) following their uptake at the axonal terminals (Figure 6). The motion characteristics of loaded vesicles with nTg and Tg dendriplexes were characterized by meticulously monitoring and tracking their trafficking along the axons (Figure 6A, 6A1 and S11, SI). Several parameters of the vesicle motion were determined, including the probability density function of instantaneous velocity, the average velocity, the recorded run length of the NPs, and the time paused.

**Figure 6.**
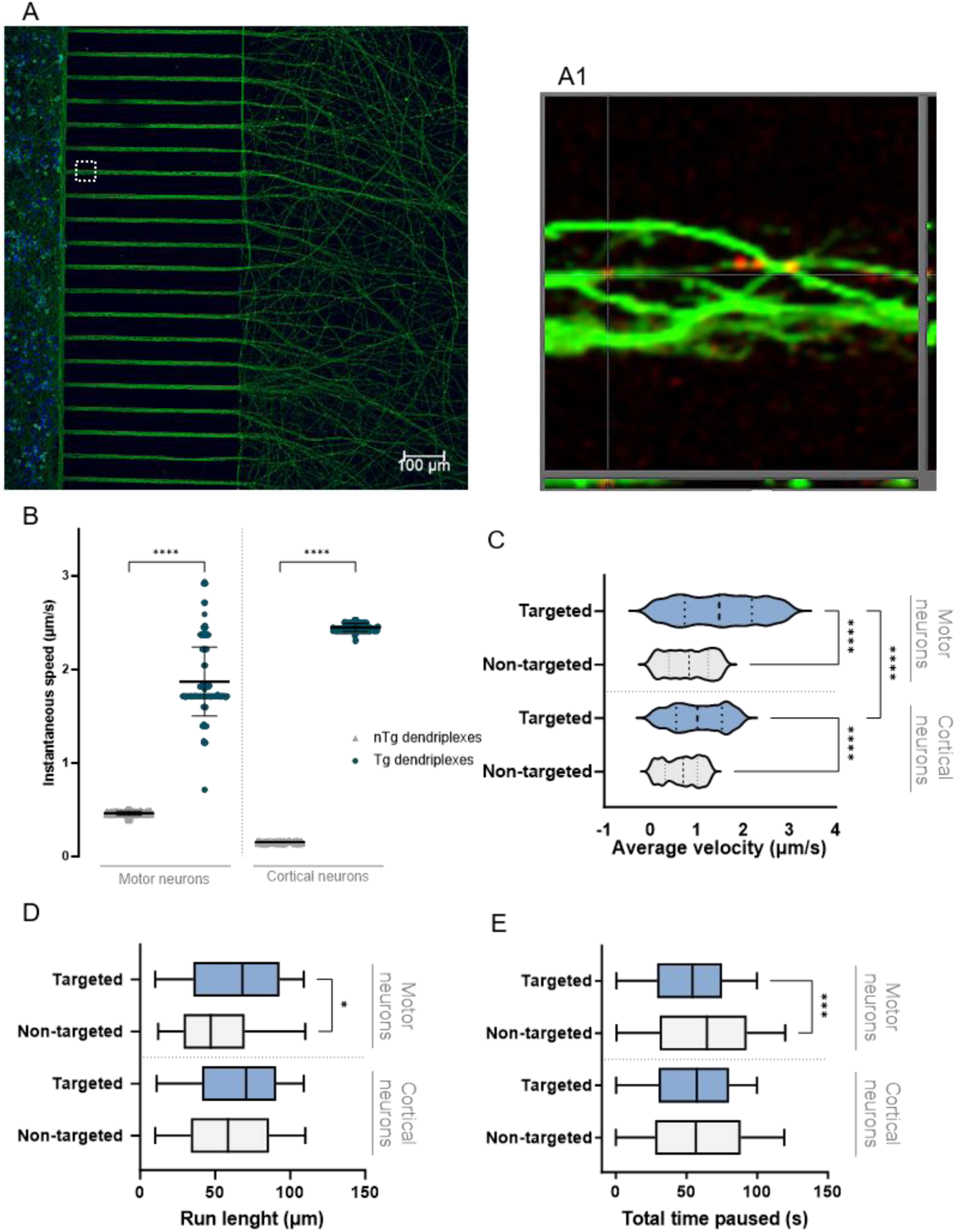
Retrograde axonal transport characteristics in motor and cortical neurons. A) Representative confocal image of cortical neurons seeded in two-compartment microchannels; (A1) Orthogonal view (XY, XZ, YZ) through axons in a microchannel (highlighted with a white dashed line in (A)) showing a nTg N/P 5 dendriplexes-loaded vesicle (red) being trafficked inside the axon, immunostained for βIII tubulin (green); (B) Instantaneous velocity frequency of nTg and Tg dendriplexes-loaded vesicles, defined as the speed (μm/s) of movement between consecutive images within a sequence. Each value represents the velocity of a single step between two frames; (C) Average velocity (μm/s) of retrograde axonal transport of NPs-loaded vesicles; (D) total run length and (E) total pause periods. Results are represented as mean ± SD of three independent experiments (n = 3), 1 replicate per experiment. Two-way ANOVA tests were used for statistical analysis. Significant differences: *p < 0.05, *** p < 0.001 and ****p ≤ 0.0001.

In both types of tested neuronal cells, the mean instantaneous velocity was significantly higher for vesicles loaded with Tg dendriplexes when compared to vesicles loaded with nTg dendriplexes (Figure 6B). In motor neurons, the mean instantaneous velocity of Tg dendriplexes was 3.9 times higher than that of nTg systems. And in cortical neurons, the difference increased to 15.3 times higher velocity. In addition, in cortical neurons, there was a much narrower distribution of velocities in the case of Tg NP-loaded vesicles. The instantaneous velocity distribution closely resembled that described for the isolated HC domain [41]. This domain was attached to the dendriplexes, which enhanced their migratory capability along the neurons, reaching the cell soma quickly. Interestingly, even though the Tg dendriplexes are functionalized with the binding domain of TeNT, which is known to primarily target peripheral motor neurons and sensory neurons [36, 42], a significantly higher instantaneous speed was observed in central cortical neurons. This reinforced the potential of Tg dendriplexes to be used in therapies targeting various neurological diseases affecting PNS and/or CNS.

Previously, we explored TMC-based NPs functionalized with the HC domain and these Tg systems exhibited both retrograde and anterograde movement. But it is worth noting that this migratory behaviour was observed in digested rat dorsal root ganglion (DRG) neurons in microfluidic culture [31]. This bidirectional movement in both the anterograde and retrograde directions is a characteristic feature of the HC domain of TeNT [41]. Here, the exclusively positive instantaneous velocities demonstrated that both Tg and nTg dendriplexes exhibited movement only in the retrograde direction along all axons of imaged embryonic mice motor and cortical neurons. With the addition of HC at the periphery of dendriplexes, they retained the ability to migrate in a unidirectional manner, and at a faster pace.

Similarly, the average velocities of the Tg dendriplexes were higher than those of the nTg dendriplexes, both in motor neurons and cortical neurons (Figure 6C). In motor neurons, the average velocity of Tg dendriplexes was around 1.48 µm/s, while in cortical neurons, the average velocity was approximately 1.01 µm/s, both in the range of fast axonal transport [31, 43]. It is important to point out that the average velocities of both nTg and Tg dendriplexes closely align with the velocities of the microtubule-dependent molecular motor dynein, which is responsible for retrograde cargo transport [44, 45]. This observation reinforces what was previously mentioned: surface functionalization of NPs with the HC fragment boosts the retrograde axonal transport capacity and migratory velocity of these NPs.

Concerning the total distance progressed per run, the Tg dendriplexes covered a greater distance, which was statistically significant in the case of motor neurons (Figure 6D). In cortical neurons, Tg and nTg dendriplexes made longer runs. On average, the Tg dendriplexes migrated 70.5 µm per run along the axons of motor neurons and 68 µm per run in cortical neurons. In motor neurons, the distance covered by the Tg systems was 1.45 times longer than by the nTg ones (nT average distance ran: 48.6 µm), while in the cortical neurons, the distance covered by the Tg systems was 1.21 times greater (nT average distance ran: 56.2 µm). Although in both types of neurons, the nTg dendriplexes covered a substantial distance, the HC endowed the dendriplexes with the ability to move over even longer distances. Finally, when evaluating the total time of NPs paused (without movement along axons), the Tg dendriplexes exhibited a significantly lower stationary time than the nTg dendriplexes in motor neurons (Figure 6E). In these neurons, the Tg systems stopped for a total of 54.2 seconds, while the nTg ones paused for a total average of 64.5 seconds. These results could be anticipated, as the average velocities of Tg dendriplexes are significantly higher, indicating that they also pause less frequently than the nTg dendriplexes. Curiously, there were not many differences in total pause times in cortical neurons. In these neurons, the nTg and Tg systems displayed similar and nearly identical pause times, 57.3 seconds and 56.5 seconds for nTg and Tg dendriplexes, respectively. Despite similar pause times, the average speed is significantly higher since when Tg dendriplexes move, they do so with a much greater instantaneous speed than nTg formulation (Figure 6B). Overall, functionalization with HC has allowed dendriplexes to move with fewer interruptions. This means that HC may have facilitated increased and/or more frequent interactions with microtubule motors.

Furthermore, the results obtained also shed light on the entry mechanisms of the nTg and Tg formulations. We have previously shown that nTg siGFP-dendriplexes internalize through a caveolin-based mechanism in mouse cortical neurons [28]. In contrast, Tg dendriplexes are expected to adopt the internalization mechanism and axonal movement of TeNT. This entails interacting with cells using a dual-receptor mechanism involving gangliosides and synaptic proteins in the neuronal membrane [46, 47], followed by internalization through a specialized clathrin-dependent pathway [48]. In fact, our results suggest that it is likely for Tg NPs to follow this internalization route, gaining access to the retrograde transport mechanism along the axon.

To evaluate the capacity to reach the cell soma of neurons, dendriplexes, prepared at N/P ratio 5 and carrying a Cy5-siRNAmi, were added in the axonal terminals compartment. Neurons were then incubated with NPs for 8, 12, 24, 48, and 120 hours, fixed, and then analysed by confocal microscopy. The Cy5 fluorescence signal was subsequently quantified in the cell soma compartment (Figure 7A).

**Figure 7.**
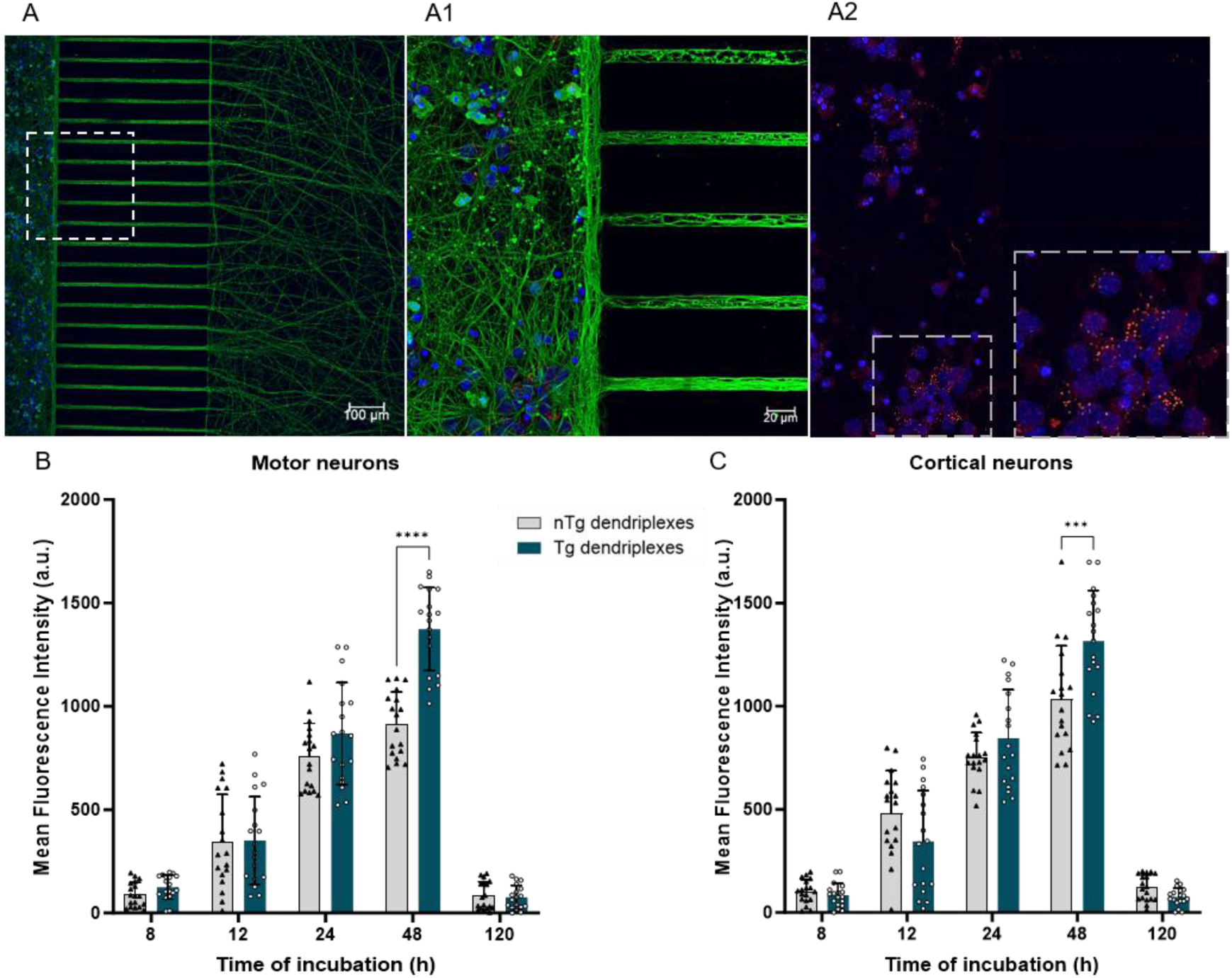
Dendriplexes’ cellular interaction kinetics assessed in microfluidic-based cultures. (A) Representative images from confocal microscopy of nTg N/P 5 dendriplexes carrying Cy5-siRNAmi in the cell soma of cortical neurons after 24 hours of incubation. (A1) and (A2) Image zoomed in on the cell soma compartment, providing a clearer identification of the dendriplexes. Staining: Nuclei with Hoechst 33 342 (in blue), βIII tubulin (in green) and Cy5-siRNAmi dendriplexes (red). Through image analysis, the fluorescence intensity in the cell bodies of (B) motor neurons and (C) cortical neurons was quantified. Results are represented as mean ± SD of three independent experiments (n = 3), with 6 replicates per experiment. For statistical analysis, the two-way ANOVA test was used. Significant differences: ***p ≤ 0.001 and ****p ≤ 0.0001.

Both motor and cortical neurons exhibited detectable Cy5 fluorescence after 8 hours of incubation with both types of dendriplexes (Figure 7B and C). As the incubation time increased, the fluorescence signal gradually and continuously intensified, reaching its peak at the 48-hour incubation time. Tg dendriplexes exhibited a higher tendency to reach and accumulate in the neuronal cell body, with a smaller quantity remaining in the axonal compartment (Figure S12, SI). As commented, notably, at 48 hours of incubation, the accumulation was more evident and statistically significant. At this time point, the cell bodies of both motor and cortical neurons showed 1.5-fold and 1.3-fold greater intensity of Cy5 fluorescence, respectively, compared to neurons that were incubated with nTg dendriplexes. This significant increase in NP presence within the cell body can be also attributed to their increased association with the neuronal cell membrane (Section 3.3.), their faster migration speed along the axon, and shorter pause times (Figure 6). This heightened presence of Tg dendriplexes in the cell soma of neurons can also account for the enhanced outcomes observed in GFP downregulation (Section 3.3). Also, to note that beyond the fluorescence intensity peak at 48 hours, a decline in the Cy5 signal intensity at the 120-hour time was observed. This decrease could potentially be attributed to vesicle dissociation.

### 3.4. Biocompatible profile of dendriplexes in microfluidic neuronal culture

To assess cellular viability following 48 hours of exposure to Tg and nTg dendriplexes (N/P 5), an LDH cytotoxicity assay in the cell soma compartment (Figure S7, SI).

This evaluation was carried out in primary motor neuron and cortical neuron cultures, which were seeded in the microfluidic devices. To understand the differences regarding cell culture conditions (conventional *versus* microfluidic), the administration of NPs to cell bodies compared to axonal terminals in a microfluidic setup was assessed. This microfluidic model enables NPs to interact with neuronal axons in a manner closely resembling *in vivo* conditions.

Our NPs, whether Tg or nTg, exhibited a low effect on the cell membrane (Figure S7, SI). This conclusion is based on the minimal percentage of LDH release by embryonic motor neurons (Figure S7A, SI) and cortical neurons (Figure S7B, SI) in microfluidic devices. Distinctly, both Tg and nTg dendriplexes showed significant differences when compared to the standard for transfections in plate assays - L2k. This lipid vector had a significant impact on the PNS and CNS neuron membranes. Notably, the biocompatibility of our vectors holds great promise, not only for *in vitro* models but also for *in vivo* applications. Also, as expected, Triton X-100 (a cell-lytic positive control) resulted in cell death and a notable release of LDH.

Overall, cells cultured in microfluidics and treated with Tg and nTg dendriplexes in the axonal compartment had a very low LDH release, similar to untreated cells. This result can be due to the lower number of NPs that reach the cell soma than the number of NPs that contact the cell soma in well-plates. This difference in NP binding can be observed in the fluorescence intensity of cells after incubation with Cy5-siRNAmi-dendriplexes in both well-plate settings (Figure 4B) and microfluidic devices (Figure 7B and 7C). The outcome observed in the microfluidic platforms holds remarkable significance, as this model reproduces the physiological separation between the cell soma and axonal terminals. This microfluidic model enables NPs to interact with neuronal axons in a manner closely resembling *in vivo* conditions [49, 50]. Aligning with our goal for peripheral administration of nanosystems, NPs establish their initial contact not with the cell soma of neuronal cells, but instead with their terminals [31]. Additionally, cells cultured in microfluidic devices and treated with NPs in the cell soma compartment showed LDH release levels similar to those observed in the well plates (data not shown), where the entire cells are exposed. Thus, it is confirmed that the mode of administration of dendriplexes has an impact on the cellular response. Interestingly, this result resembles what we previously obtained in a study of TMC-based NPs in microfluidic DRG cultures [31].

To evaluate the biocompatibility through the measurement of the neuronal electrical activity, neurons were incubated in three-compartment microfluidic cultures, in which the proper alignment of the MEAs with the compartments and microchannels was confirmed (Figure 8A and B). After incubation with our NPs, the electrical activity of neurons was measured (Figure 8C and S13, SI). No electrophysiological alterations were identified in either motor neurons or cortical neurons following incubation with Tg or nTg N/P 5 dendriplexes. The total number of spontaneous spikes and other measures of neuronal electrical activity (number of bursts, ISI, and IBI) were similar to those in the untreated group. This demonstrates that the addition of NPs to the axons did not affect neuronal viability or activity. Furthermore, the spiking data was comparable between samples treated with NPs and untreated samples (FigureS13M, SI). This shows that the addition of NPs to the axons of motor neurons did not affect the viability and activity of the co-culture, and notably, it had no impact on central neurons electrophysiological performance. This electrophysiological response further strengthens the neuro-applicability of the developed vectors.

**Figure 8.**
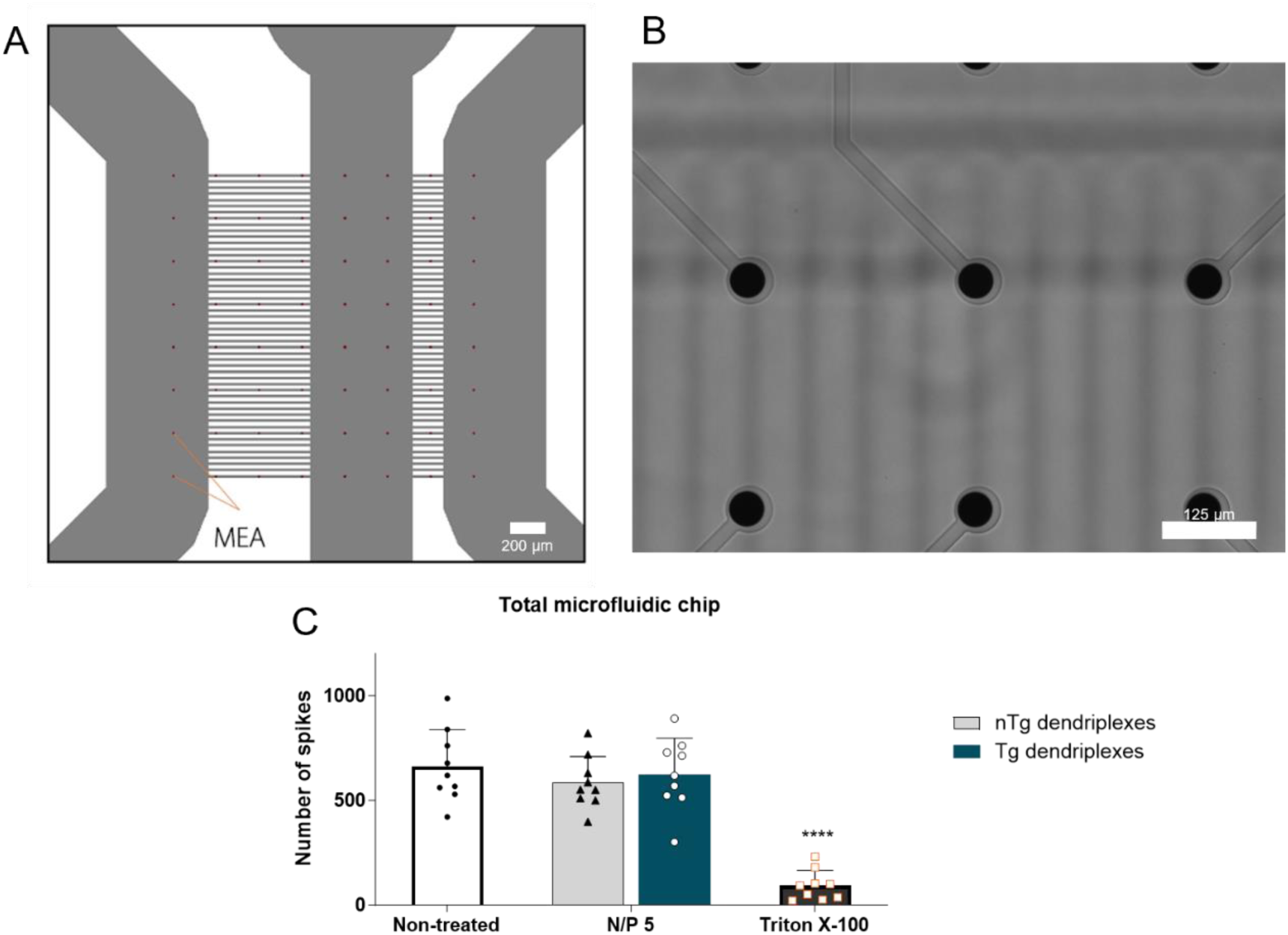
Biocompatible profile evaluation in primary motor and cortical neurons. (A) Computational prediction of the alignment of MEAs and PNS-CNS-on-Chip; (B) Microscope image confirmed the right alignment of MEAs and microchannels (scale bar: 125 µm). The electrical activity was also measured for 5 minutes along (C) all microfluidic chips. Results are represented as mean ± SD of three independent experiments (n = 3) with 3 microfluidics per experiment. For statistical analysis, the two-way ANOVA test was used. Significant differences: *p < 0.05, **p < 0.01, *** p < 0.001 and ****p ≤ 0.0001.

### 3.5. The enhanced axonal outgrowth after dendriplexes treatment

The proposed NPs were further assessed for their capacity to carry and release siPTEN, resulting in the downregulation of a biologically relevant gene (PTEN) through axonal outgrowth evaluation.

The culture of embryonic cortical neurons was conducted within three-compartment microfluidic platforms featuring two sets of microchannels, each measuring 600 µm in length (Figure 1B). This setup enabled us to have two distinct axonal compartments for evaluation. Neurons were seeded in the central compartment, and within this same compartment, Tg and nTg dendriplexes carrying siPTEN were added, enabling us to scrutinize the changes in axon development resulting from PTEN silencing. Five days after the NPs addition, the cultures were fixed, and axonal growth was assessed through image analysis. Linear Sholl analysis MATLAB scripts were applied to analyse axonal growth within the axonal compartments (Figure 9A) [35]. When neurons were treated with nTg or Tg dendriplexes, a higher number of intersections per grid line normalized to the number of neurons were recorded in relation to untreated neurons (Figure 9B). This implies that a greater number of axons traverse the microchannels, reaching longer distances of up to 800 µm. The average length of axons when treated with nTg and Tg dendriplexes was significantly increased, measuring 194 and 256 µm, respectively (Figure 9C). These lengths were 1.3 and 1.7 times greater than the average axon length of untreated cells, indicating a notable enhancement in axonal growth. Interestingly, dendriplexes prepared at an N/P 5, especially Tg dendriplexes, exhibited a more favourable axonal response compared to those at an N/P ratio of 10. This may be attributed to the fact that N/P 5 dendriplexes contain a lower amount of fbB dendrimer, so they are less stable, facilitating the release of siPTEN, as seen in the results after incubation with heparin (Section 3.1.).

**Figure 9.**
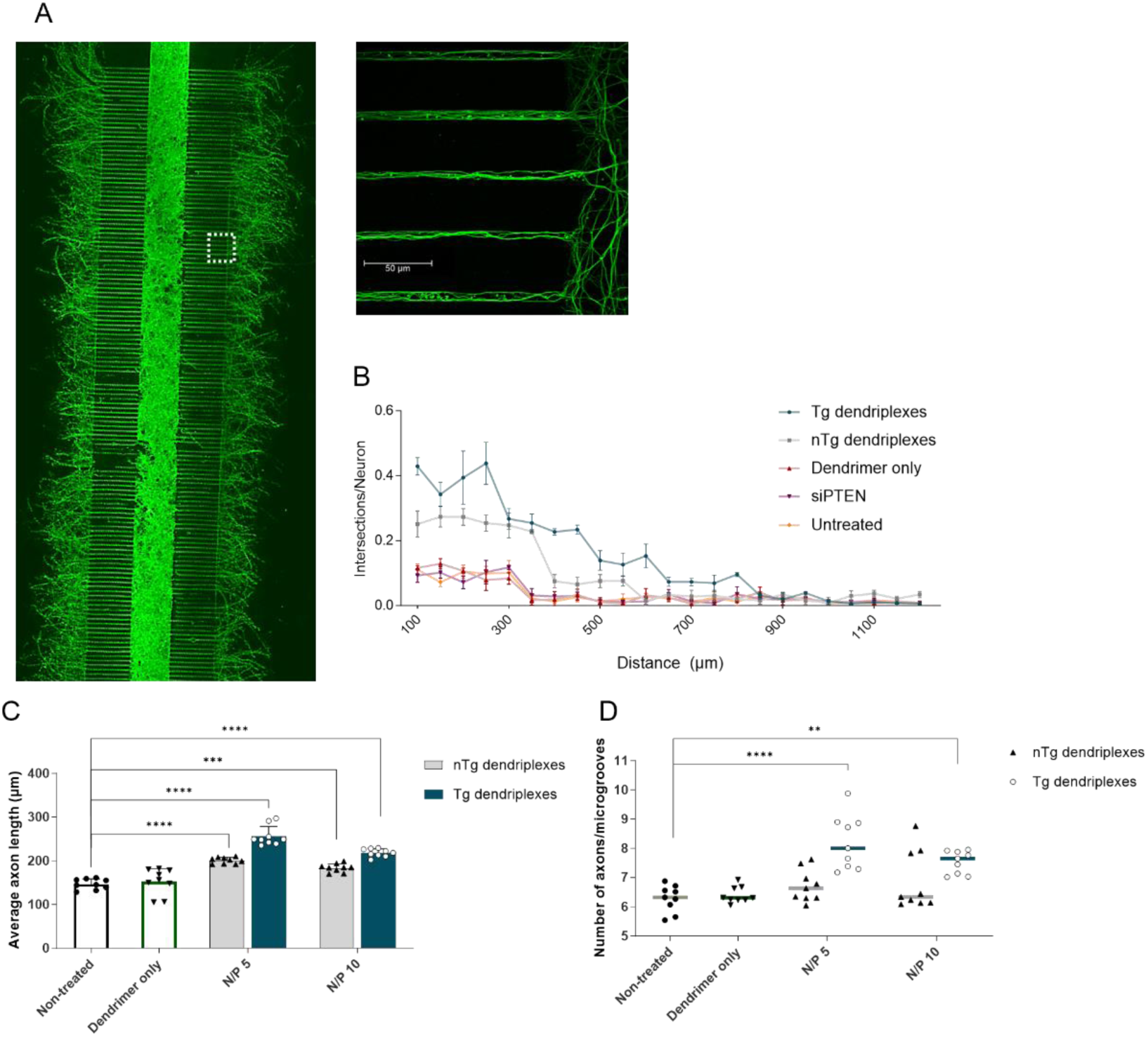
Biological effect of our N/P 5 or 10 NPs formulations in cortical neurons. Representative mosaic image used to quantify axonal outgrowth, in three-compartment microfluidic devices, after treatment with targeted N/P 5 dendriplexes carrying siPTEN. Staining: Neurofilament H (in green); (B) number of intersections of the axons in linear grids spaced 50 µm apart per neuron after incubation with N/P 5 dendriplexes; (C) average axon length and (D) number of axons per microchannels. Results are represented as mean ± SD of three independent experiments (n = 3), 3 replicates per experiment. For statistical analysis, the two-way ANOVA test was used. Significant differences: *p < 0.05, *** p < 0.001 and ****p ≤ 0.0001. In (B), the statistical analysis was performed in comparison to the untreated samples.

Furthermore, Tg dendriplexes demonstrated a substantial advantage in promoting axonal outgrowth (Figure 9D). The treatment with Tg dendriplexes led to a significantly increased number of axons that traverse the microchannels and reach the axonal compartment, with approximately 8 axons observed when treated with N/P 5 Tg dendriplexes and 7.7 when incubated with N/P 10 Tg dendriplexes. In comparison to untreated cells (approximately 6.3 axons per microgroove), the axon count increased by approximately 1.3 and 1.2-fold for Tg N/P 5 and 10 dendriplexes, respectively. This augmentation in axon quantity may also be attributed to PTEN silencing.

Altogether, this increased axonal outgrowth and the boost in axon length underscore the potential of Tg dendriplexes for enhancing axonal development.

### 3.6. Targeted dendriplexes transcytosis in PNS-CNS-on-Chip

In addition to their potential application in both PNS and CNS neurons, the capacity of Tg dendriplexes to traverse the PNS and reach the CNS was sought. To evaluate this, we used the three-compartment microfluidic device to develop a PNS-CNS-on-Chip system (Figure 1C, 1D and 10A). In this case, the microchannels had different lengths - ones measuring 450 µm and the others 200 µm in length. These two different sets of microchannels were designed to replicate the smaller length of axons of CNS neurons and the greater axonal length of peripheral neurons.

In this platform, we seeded motor neurons in the central compartment and facilitated axonal growth within the 450 µm microchannels. Twenty-four hours later, cortical neurons were seeded in the lateral compartment and the axonal growth within the 200 µm microchannels was promoted by hydrostatic pressure. On DIV5 after motor neuron seeding, dendriplexes at N/P 5, prepared with Cy5-siRNAmi (final concentration of 100 nM), were added into the free lateral (peripheral) compartment (Figure 1C and D, compartment 3). After 72, 96, and 120 hours, the cells were fixed and subjected to microscopic evaluation. In the first two tested time points, no Cy5 signal was observed in cortical neurons, only in motor neurons, with a slight accumulation of the signal in these neurons. After 120 hours of incubation, the Cy5 fluorescence signal was clearly identified in motor neurons (Figure 10B), and it was also detected in the cell bodies of cortical neurons (Figure 10C). Interestingly, only the Tg dendriplexes could reach the CNS compartment (Figure 10C, D and E). Furthermore, notably, the Cy5 signal identified in neurons at 120 hours differs from that observed in neurons after 24 hours (Figure 7A). After 24 hours of incubation, the Cy5 signal appears more punctate, possibly NPs are still within vesicles. After 120 hours of incubation, the signal appears dispersed, which could be caused by the release of dendriplexes and/or nucleic acid (Figure 10B and C) [51].

**Figure 10.**
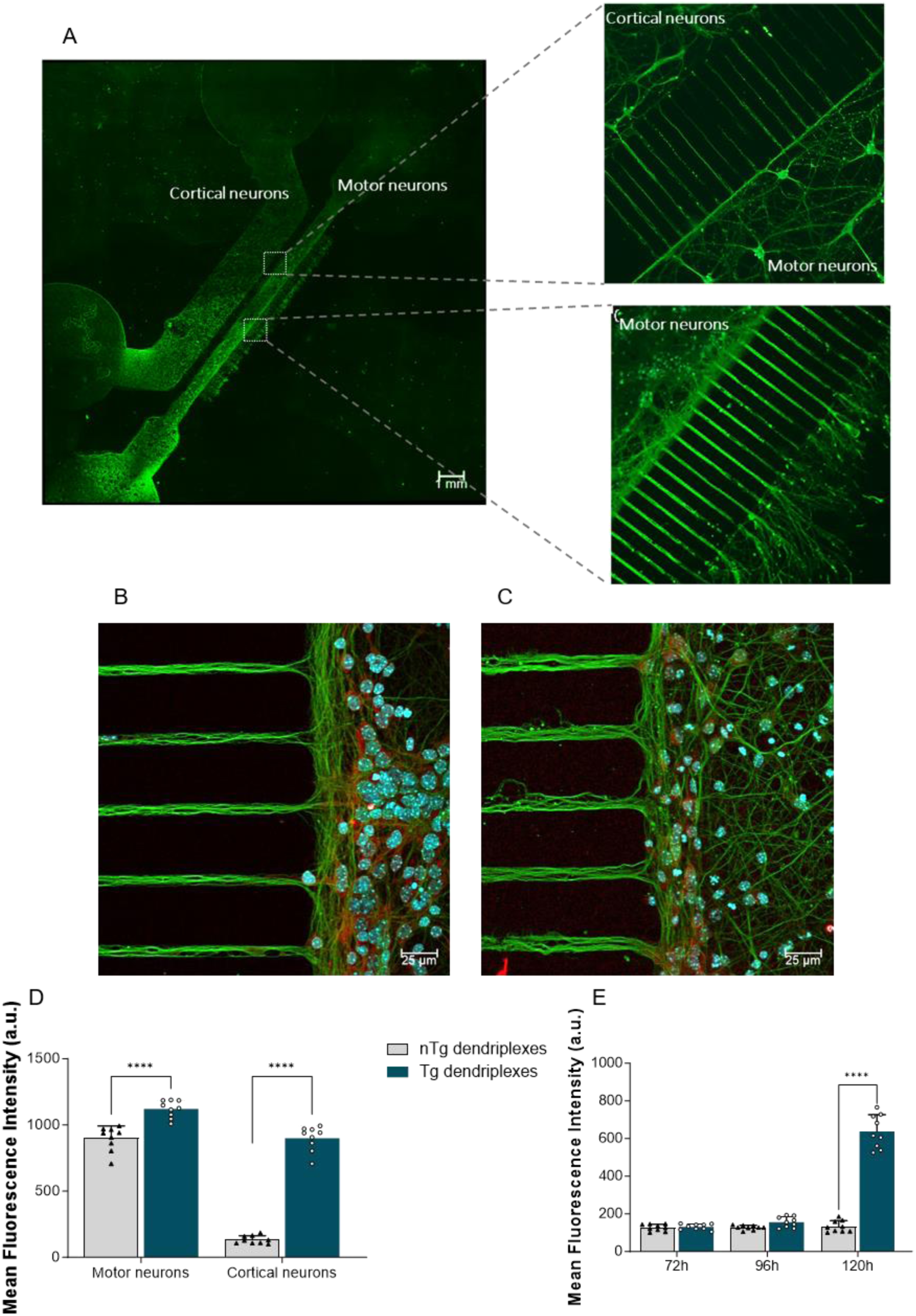
Transcytosis capacity in PNS-CNS-on-Chip. (A) Representative images of widefield microscopy of the developed PNS-CNS-on-Chip; Images of (B) motor neurons and (C) cortical neurons of the PNS-CNS-on-Chip after incubation with nTg and Tg dendriplexes for 120 hours. Staining: Nuclei (in cyan), βIII tubulin (in green) and Cy5-siRNAmi dendriplexes (red). Through image analysis, (D) mean Cy5 fluorescence intensity was quantified in the motor neurons compartment and cortical neurons compartment after 120 hours of incubation with NPs. (E) Kinetics of internalization in cortical neurons at 72, 96, and 120 hours. Results are represented as the mean of three independent experiments (n = 3), 3 replicates per experiment. For statistical analysis, the one-way ANOVA test was used. Significant differences: *p < 0.05, *** p < 0.001 and ****p ≤ 0.0001.

Beyond the biocompatible nature of our dendritic NPs in PNS and CNS neurons and their capacity to be transported inside neurons, their ability to transcytose between neurons enhances the potential for treating neurological disorders, both in the PNS and CNS. Our Tg dendriplexes demonstrated the ability to migrate to CNS neurons through PNS neurons, and to the best of our knowledge, this is the first time such capability has been demonstrated by NPs.

### Conclusions

Our study focused on developing, characterizing, and exploring the neuro-applicability of neuron-Tg fbB-based dendriplexes, functionalized with the neurotropic TeNT HC domain, for siRNA delivery. Exploring the interaction between fbB dendrimer and siRNA effective siRNA complexation was demonstrated across different N/P ratios, unaffected by TeNT protein fragment functionalization. Evaluation of assembly reversibility revealed substantial siPTEN release, highlighting the vector’s payload-delivering proficiency. Furthermore, dendriplexes showed spherical morphologies and excellent nanosizes conducive for neuronal applications, consistently below 200 nm, with narrow PDIs.

The LDH cytotoxicity assay showed minimal impact on cell membranes, indicating the safety of both nTg and Tg NPs in traditional neuronal cultures. Neurotropism studies confirmed the superior affinity of Tg dendriplexes for both PNS and CNS neurons, emphasizing their efficient targeting with the TeNT protein fragment. The downregulation of GFP expression further highlighted the promising potential of Tg dendriplexes as effective and safe neuronal siRNA delivery vectors.

In microfluidic-based neuronal cultures, Tg dendriplexes demonstrated robust performance. Our microfluidic chips mimicked natural neural architecture, allowing precise assessment of NP-cell interactions. Live cell imaging showed enhanced retrograde transport of Tg dendriplexes. Moreover, both nTg and Tg dendriplexes exhibited exceptional biocompatibility in microfluidic cultures and electrical activity analysis confirmed their neuro-applicability without compromising healthy neurons.

The biological impact of dendritic NPs was assessed in embryonic cortical neurons within three-compartment microfluidic platforms. The analysis revealed enhanced axonal proliferation and length, with Tg dendriplexes showing a significant advantage, indicating their potential for promoting axonal development.

Lastly, the transcytosis capability of Tg dendriplexes was demonstrated in a PNS-CNS-on-Chip microfluidic system, showcasing successful migration from the motor to cortical neurons. The distinct Cy5 signal over time suggested efficient transport, highlighting the therapeutic potential of Tg dendriplexes for neurological disorders in both PNS and CNS.

Altogether the results underscore the great potential of our fully biodegradable Tg fbB-based dendriplexes for effective siRNA delivery in both PNS and CNS neurons, opening new avenues for research in the Neurosciences and gene therapy fields.

## Conflicts of interest

The authors assert that there are no potential conflicts of interest that could have had any influence on the work presented in this paper.

## Acknowledgements

This work was supported by Portuguese funds through *Fundação para a Ciência e a Tecnologia*, I. P. (FCT) in the framework of the projects PTDC/CTM-NAN/3547/2014 and PTDC/BTM-MAT/4156/2021. A. P. S. acknowledges FCT for her Ph.D. scholarship (SFRH/BD/137073/2018), EMBO for her Scientific Exchange Grant (10398) and COST for her short-term scientific mission grant (9f51e2a4, COST Action: CA17103 - Delivery ofAntisense RNA ThERapeutics (DARTER) Action). V. L. appreciates her Assistant Researcher contract under the “Concurso Estímulo ao Emprego Científico Individual – 4.ª Edição” (2021.00472.CEECIND), and S. C. G. for her post-doctoral fellowship (SFRH/BPD/122920/2016). B. M. M. also acknowledges the support of the Israel Science Foundation (ISF grant: 2248/19, 1934/23), ERC SweetBrain 851765, The Aufzien Family Center for the Prevention and Treatment of Parkinson’s Disease at Tel Aviv University, The Zimin Foundation, the Innovation Authority “BioChip” and the Israel Ministry of Science and Technology (Grant No. 3–17351, 8-2175), the European Union Horizon 2020 research and innovation programme under the Marie Skłodowska-Curie (grant agreement No. 101007804-Micro4Nano) and a OSTEONET project grant (agreement No 101086329). The size and zeta potential analyses were conducted at the Biointerfaces and Nanotechnology Scientific Platform, with the assistance of Ricardo Vidal. Confocal microscopy was conducted at the i3S Scientific Platforms Bioimaging and at Advanced Light Microscopy, at the latter with the assistance of Dr Maria Azevedo and Dr Paula Sampaio. These platforms are members of the Portuguese Platform of Bioimaging (PPBI; PPBI-POCI-01-0145-FEDER-022122). The authors acknowledge Ana Castanheira at the INL for her support in using the Nanosight.

## Notes

### Competing Interest Statement

The authors have declared no competing interest.

