## Supporting Information for "Unveiling the potential of neuron-targeted dendriplexes for siRNA delivery using a PNS-CNS-on-Chip"

### Table of Contents

Page


1) siRNA sequences 3
2) Development of microfluidic platforms to test the intra- and intercellular pathways of dendritic NPs 4
3) siRNA release capacity of non-targeted dendriplexes 9
4) Biocompatibility evaluation in well-plates and microfluidic devices 10
5) Microfluidic platforms characterization 13
6) Axonal transport of neuron-targeted dendriplexes 16
7) Evaluation of the capacity to reach cell body of neurons – Axonal compartment 17
8) Biocompatibility evaluation in PNS-CNS-on-Chip 18

#### 1) siRNA sequences

**Table S1.** Nucleic acid sequences used in this study and their correspondent sequences.

| Nucleic acid | Sequence (5’ – 3’) |
|---|---|
| siPTEN | sense: CGACUUAGACUUGACCUAUAU<br>antisense: AUAGGUCAAGUCUAAGUCGAA |
| siGFP | sense: GCUGACCCUGAAGUUCAUCUGCACC<br>antisense: GGUGCAGAUGAACUUCAGGGUCAGCUU |
| Cy5-siRNAmi | sense: GGTGCAGATGAACTTCAGGGTCAGCTT<br>antisense: /Cy5/GCTGACCCTGAAGTTCATCTGCACC |

2’-*O*-methyl RNA bases are underlined

2) **Development of microfluidic platforms to test the intra- and intercellular pathways of dendritic NPs**

Microfabrication methods have been employed to create microfluidic platforms capable of emulating intricate three-dimensional environments, closely simulating in vivo conditions. These platforms enabled the evaluation of the intra- and intercellular transport of dendritic nanoparticles.

*Design*

Devices were meticulously designed to mimic the interface between the peripheral and central nervous systems (PNS and CNS, respectively). Three distinct device designs were engineered utilizing CleWin software (CleWin 4.0 Layout Editor). These designs feature cell culture chambers interconnected by microchannels with a 2x10-µm cross-section, with variable lengths (200 to 600 µm).

*Device fabrication – Photolithography*

Microfluidic devices were fabricated using poly(dimethylsiloxane) (PDMS) by soft lithography. Master molds were created on silicon wafers, featuring multi-height structures, employing a hybrid microfabrication technique that combined deep reactive ion etching (DRIE) and SU-8 photolithography.

To fabricate the 2-µm high microchannels, the silicon wafer initially underwent a patterning process. The wafer was vapor-primed with hexamethyldisilazane (HDMS) for 5 minutes at 150 °C to enhance resist adhesion. Subsequently, the wafer was placed in an automatic spin coater (Karl Suss Optical Track) for the application of a 2.2 µm layer of AZP4110 photoresist, spun at 1500 rpm, and soft-baked at 110 °C for 80 seconds. The microchannel design was patterned by direct write laser (DWL 2000) following which the wafer underwent chemical development for 240 seconds using a 1:4 mixture of AZ 400k developer.

The pattern integrity and the thickness of the photoresist were examined using two an OPM Nanocalc Optical Profilometer (OP) and a KLA Tencor P-16 Contact Profilometer, which indicated that the thickness of the photoresist was 2.3 µm on average.

Optical observation using a Nikon Eclipse L200N confirming the accurate replication of the design on the photoresist (Figure S1).

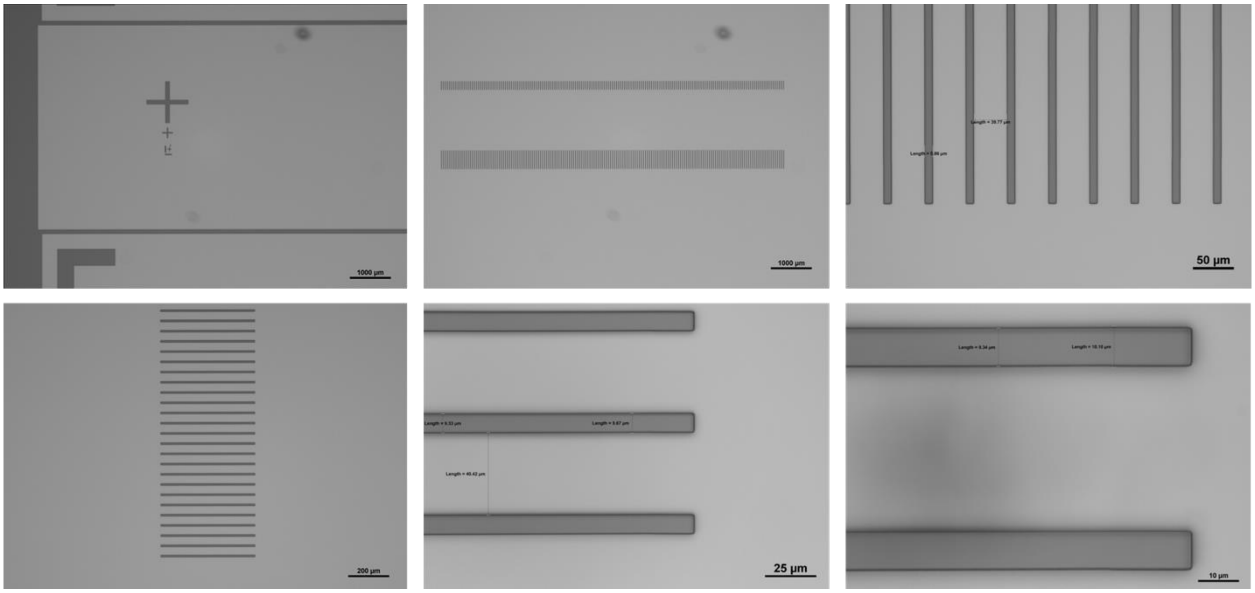


**Figure S1.** Images of microgrooves pattern (in dark grey) acquired to verify the photoresist pattern following exposure to the direct write laser (DWL).

The next step involved the direct fabrication of 2-3 µm-high microchannels on the silicon substrate through deep reactive ion etching (DRIE). A dry etching system (SPTS Pegasus) was employed, following the Bosch process. The Bosch process is a two-step procedure involving an initial isotropic silicon etch, followed by a deposition step to safeguard the sidewalls against further etching. This process was based on alternating cycles of SF6 gas for rapid silicon etching and C4F8 to create a fluorocarbon polymer deposition to passivate the sidewalls during etching.

The calculation of etch times was based on two factors: the silicon etch rate (3329 nm $min^{-1}$) and the desired target depth, which yielded an estimated etching time of approximately 1 minute and 10 seconds. Finally, the photoresist was removed using a PVA Tepla Plasma Asher, and the completed microchannel structures were inspected using a Eclipse L200N Optical Microscope (Figure S2).

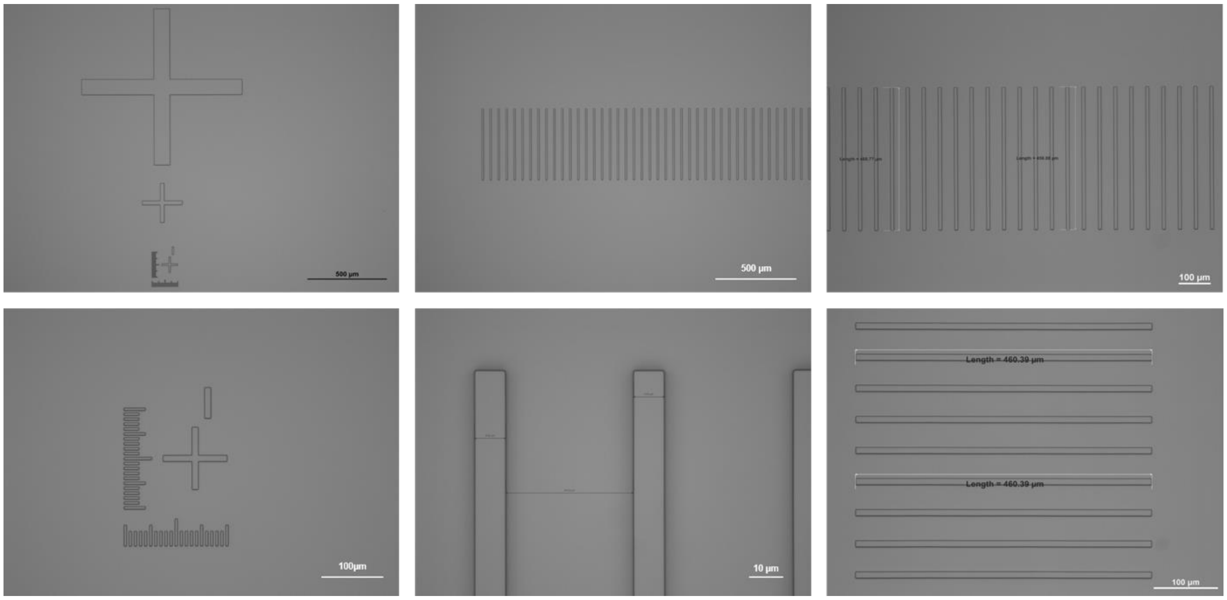


**Figure S2.** Optical microscopy images of the microchannels and alignment marks after DRIE and resist strip.

To produce the larger chambers, the wafer was cleaned by oxygen plasma cleaning followed by spin-coating with SU-8 2100, at 3000 rpm for 30 seconds. Subsequently, the wafer was placed on a hot plate for a soft bake, starting at 65 °C for 5 minutes and then ramping up to 95 °C for an additional 20 minutes. After cooling to room temperature, the wafer was exposed in a Karl Suss MA6/BA6 Mask Aligner under UV light using a chrome soda lime photomask from JD PhotoData (Herts, UK). The exposure time was 21.2 seconds to achieve in a total exposure energy of 240 mJ $cm^{-2}$. Post-exposure baking comprised an initial phase at 65 °C for 5 minutes, followed by 95 °C for 10 minutes, and a 30-minute rest. Non-exposed SU-8 was removed by immersing the mold in propylene glycol methyl ether acetate (PGMEA) solvent or SU-8 developer for 10 minutes, accompanied by gentle shaking. Finally, the wafer underwent a hard bake for 2 minutes at 150 °C.

Upon the completion of the microfabrication process (Figure S3), the feature height was confirmed using a KLA Tencor P-16 Contact Profilometer (Figure S4A), and the wafers were subjected to optical microscopy inspection (Figure S4B and C).

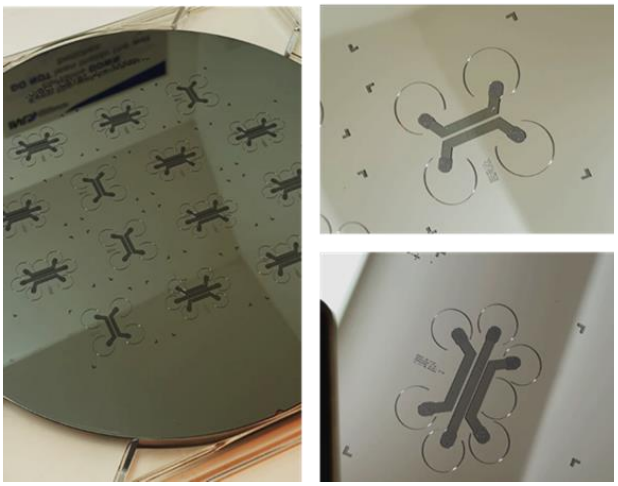

**Figure S3.** Silicon master mold after microfabrication.

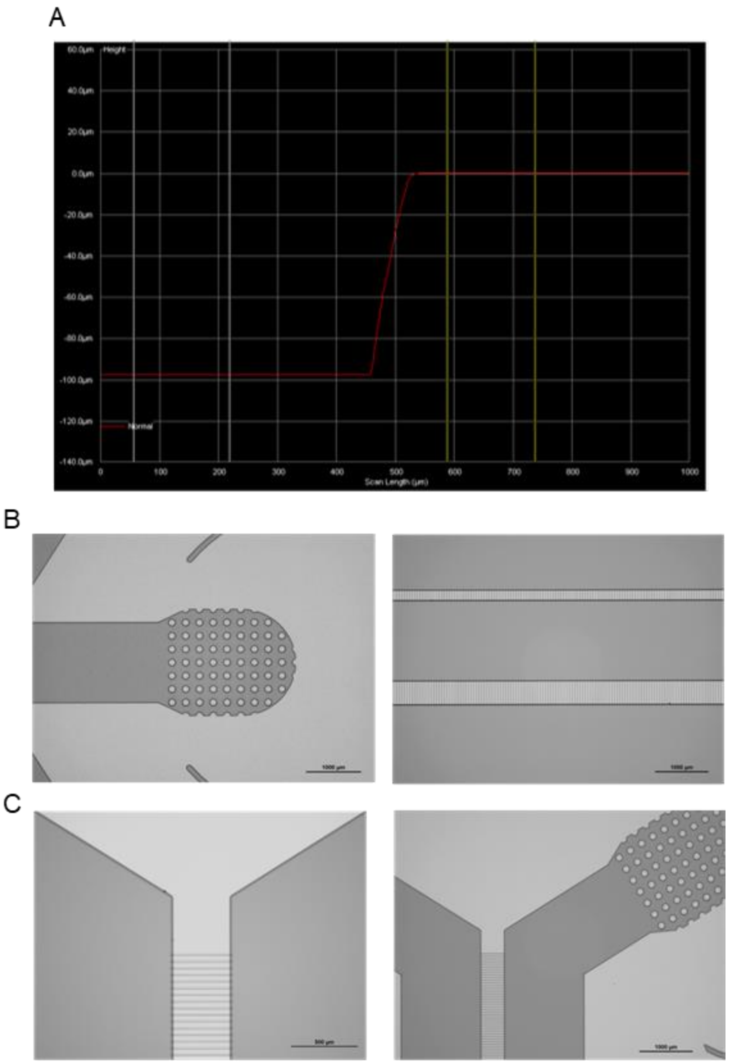


**Figure S4.** Assessment of features of microfluidic devices. (A) Representative image of profilometer measurements of the feature height of the chambers at the end of the SU-8 photolithography; (B) Optical microscope images of devices pattern of three-compartment microfluidics (microgrooves of 200 and 450 µm length); (C) Optical microscope images of devices pattern of two-compartment microfluidics (microgrooves of 450 µm length).

*Soft lithography*

PDMS was prepared by mixing the elastomer and curing agent at a 10:1 mass ratio. After thorough degassing, the mixture was poured onto the wafer (Figure S3). To define the device dimensions, an acetal frame machined by CNC micromilling was used (CNC FlexiCAM - High. The PDMS batches were then left to cure at 60 °C and demolded.

#### 3) siRNA release capacity of non-targeted dendriplexes

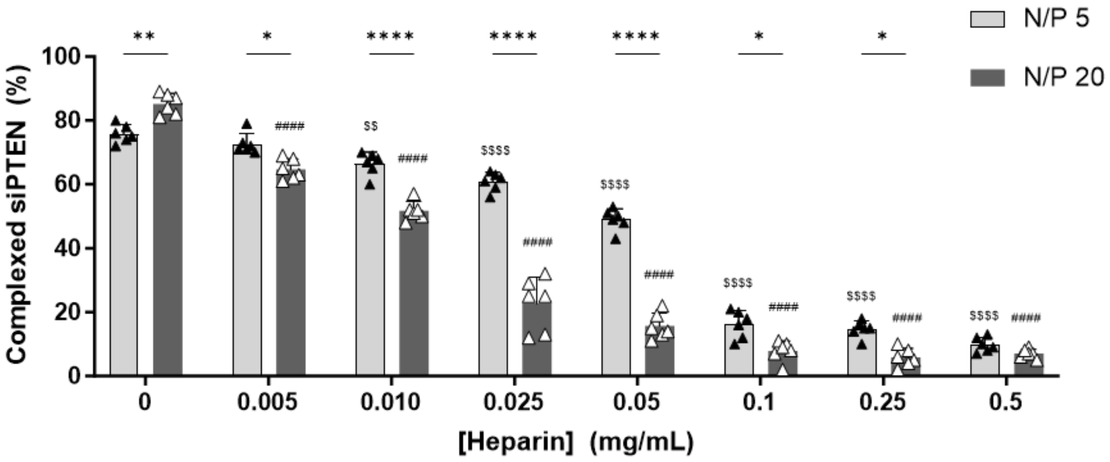


**Figure S6.** Heparin dissociation assay in non-targeted (nTg) dendriplexes prepared with siPTEN as a function of N/P ratio. Results are expressed as mean ± SD (standard deviation) of three independent experiments (n = 3), with 2 replicates per experiment. For statistical analysis, one-way ANOVA test was used. Significant differences: *p < 0.05, **p < 0.01, *** p < 0.001 and ****p ≤ 0.0001. For each concentration of heparin, the symbol $ indicates significant differences in comparison to the complexation of N/P 5 dendriplexes without incubation with heparin (heparin concentration of 0 mg/mL), while # indicates significant differences compared to the complexation of N/P 20 dendriplexes without incubation with heparin (heparin concentration of 0 mg/mL).

#### 4) Biocompatibility evaluation in well-plates and microfluidic devices

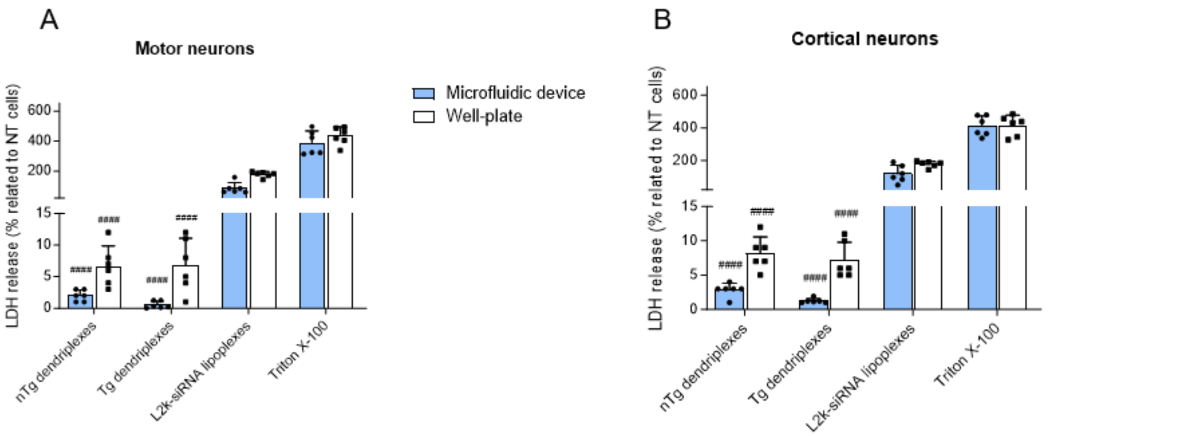


**Figure S7.** Plasma membrane integrity of (A) motor neurons and (B) cortical neurons seeded in well-plates or in microfluidic platforms, assessed by LDH assay after incubation with N/P 5 siPTEN-dendriplexes. Results are expressed as mean ± SD of three independent experiments (n = 3), with 2 replicates per experiment. For statistical analysis, one-way ANOVA test was used. For each formulation, the symbol # indicates significant differences in comparison to the LDH release values of L2k-siRNA lipoplexes and Triton X-100. Significant differences: *$p < 0.05$, **$p < 0.01$, *** $p < 0.001$ and **** or $^{\#\#\#\#}p \leq 0.0001$.

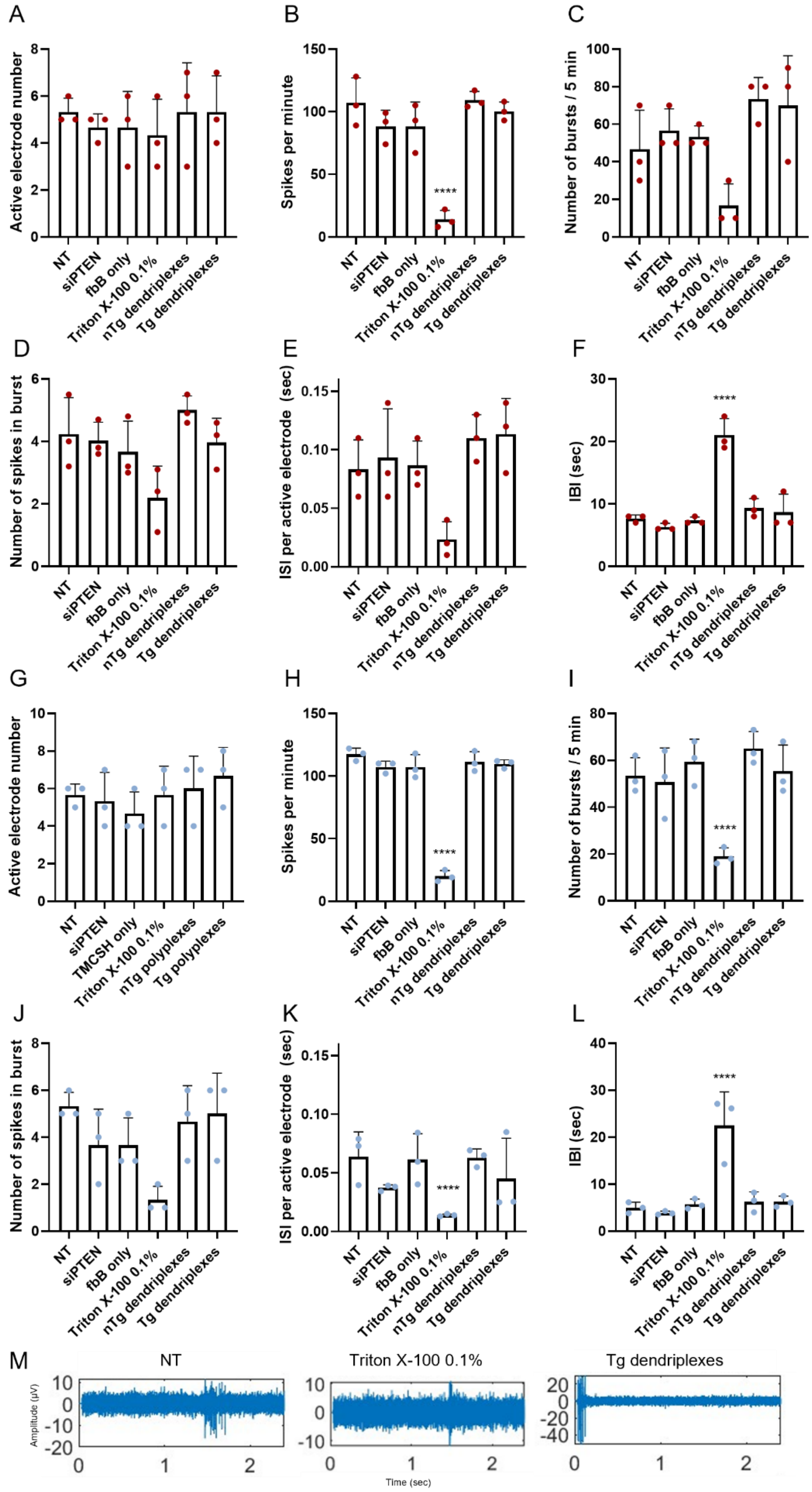

A
Active electrode number
NT
siPTEN
fbB only
Triton X-100 0.1%
nTg dendriplexes
Tg dendriplexes
B
Spikes per minute
****
C
Number of bursts / 5 min
D
Number of spikes in burst
E
ISI per active electrode (sec)
F
IBI (sec)
****
G
Active electrode number
TMCSH only
nTg polyplexes
Tg polyplexes
H
Spikes per minute
****
I
Number of bursts / 5 min
****
J
Number of spikes in burst
K
ISI per active electrode (sec)
****
L
IBI (sec)
****
M
NT
Triton X-100 0.1%
Tg dendriplexes
Amplitude (µV)
Time (sec)

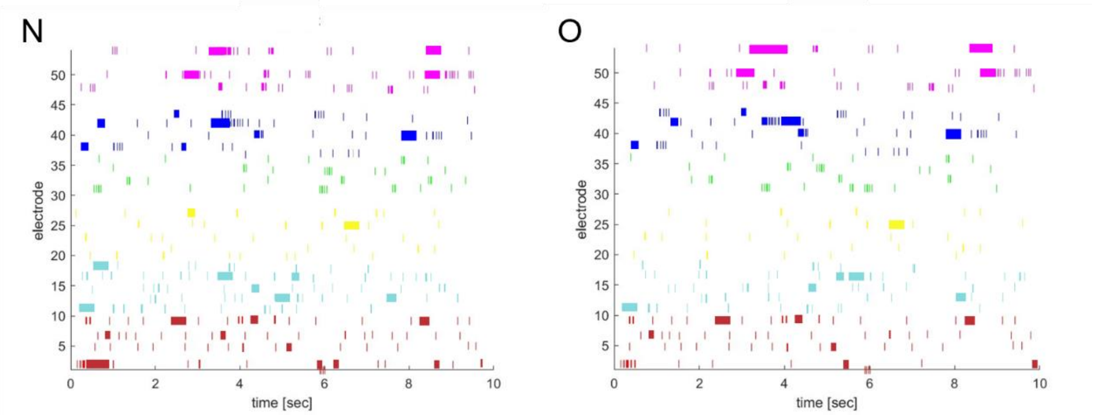


**Figure S8.** Spontaneous electrical activity of (A-F and N) motor and (G-L and O) cortical neurons cultured in well plates recorded with microelectrode arrays (MEAs) after incubation with nTg or Tg dendriplexes carrying siPTEN (DIV14). Cells treated with Triton™ X-100, 0.1% (v/v) in PBS 1× were used as controls. (A) and (G) Number of active electrodes. Active electrodes were defined by showing 15 or more spikes per minute; (B) and (H) Number of spikes per minute; (C) and (I) number of bursts during 5-min recording. Bursts were defined as three or more spikes with a 300-ms interspike interval (ISI); (D) and (J) number of spikes in burst; (E) and (K) ISI per active electrode; (F) and (L) mean intra-burst interval (IBI). Results are represented as the mean of three independent experiments (n = 3). For statistical analysis, a two-way ANOVA test was used. No statistically significant differences were identified resulting from treatment with nTg or Tg dendriplexes (N/P 5) compared to the non-treated (NT) cells. All graphs use a voltage-threshold based algorithm applied to 20,000 Hz high-pass filtered traces, with the threshold set to 4.5 times the standard deviation of the noise. (M) Representative spontaneous electrophysiological activity of NT neurons, neurons treated with Triton™ X-100, 0.1% (v/v) in PBS 1×, and neurons treated with Tg dendriplexes (N/P 5). Raster plot of spikes from more than 20 electrodes in cultures of (N) motor neurons or (O) cortical neurons, showing the pattern and density of spikes. Results are expressed as mean ± SD of three independent experiments (n = 3), with 1 replicate per experiment. For statistical analysis, two-way ANOVA test was used. Significant differences: **** $p \leq 0.0001$.

#### 5) Microfluidic platforms characterization

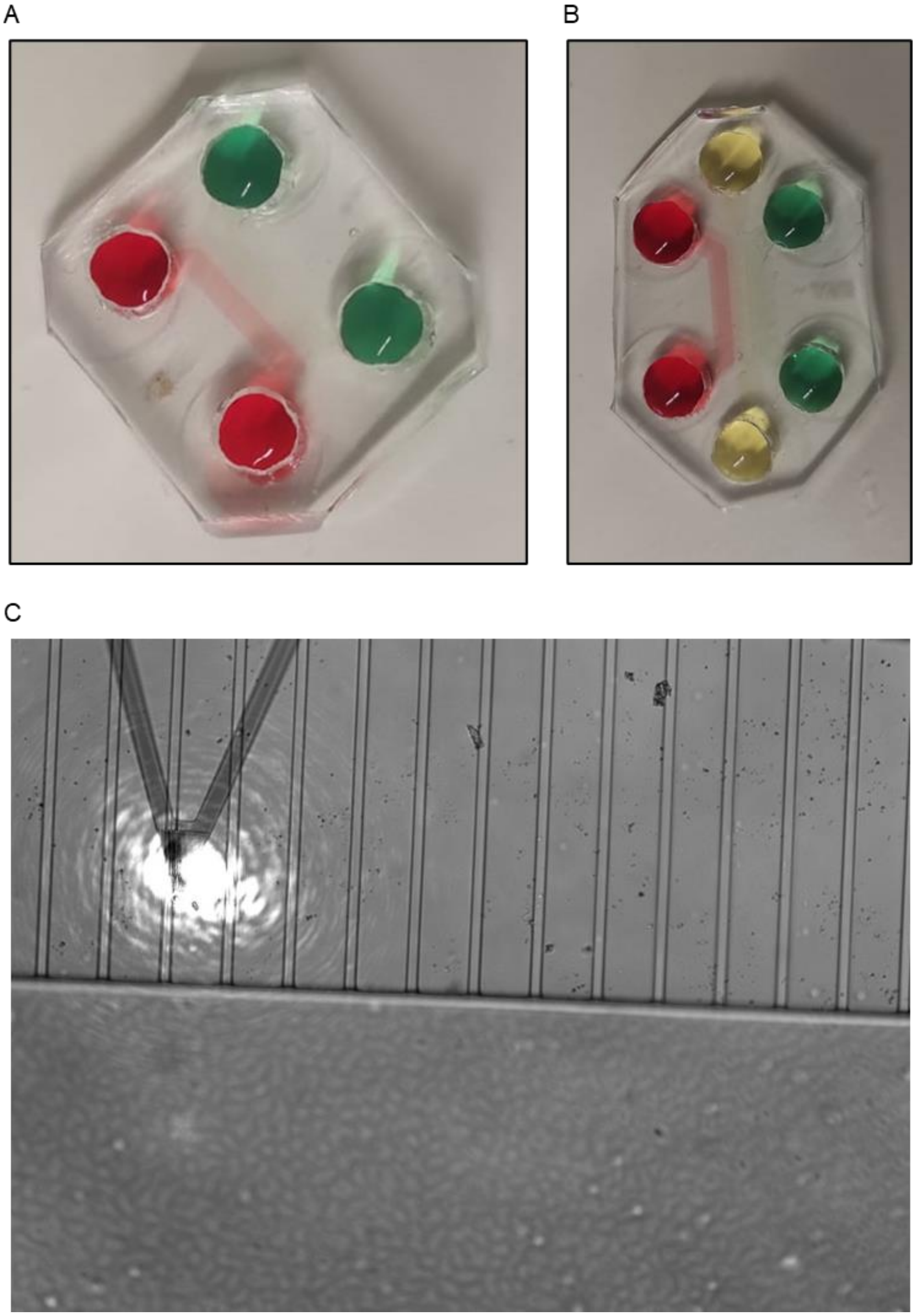


**Figure S9.** Microfluidic platforms in PDMS. (A) Microfluidics with 2 food dyes in 4 reservoirs and 2 compartments/chambers; (B) Microfluidics with 3 food dyes in 6 reservoirs and 3 compartments; (C) Representative image depicting the assessment of microchannel height using atomic force microscopy (AFM).

**Table S2.** Summary table detailing the device specifications.

| CHANNELS | Height | 100 µm |
|---|---|---|
| Microchannels | Number | 139 |
| | Length | 450 µm (two-compartment devices)<br>600/600 µm (three-compartment devices)<br>450/200 µm (PNS-CNS-on-Chip |
| | Height | ~2 µm |
| | Opening | 10 µm |

**Table S3.** Fluidic resistance of the compartment and microchannels

| Device | Fluidic resistance (Pa.s/$m^3$) |
|---|---|
| Compartment | 1.00 $x10^{11}$ |
| Microchannel | 7.72 $x10^{16}$ |

**Table S4.** Features of the inlets/outlets.

| | Diameter (m) | Area ($m^2$) | Height (m) | ΔP (Pa) |
|---|---|---|---|---|
| Inlet/outlet | 5 $x10^{-3}$ | 1.96 $x10^{-5}$ | 1.27 $x10^{-3}$ | 12.49 |

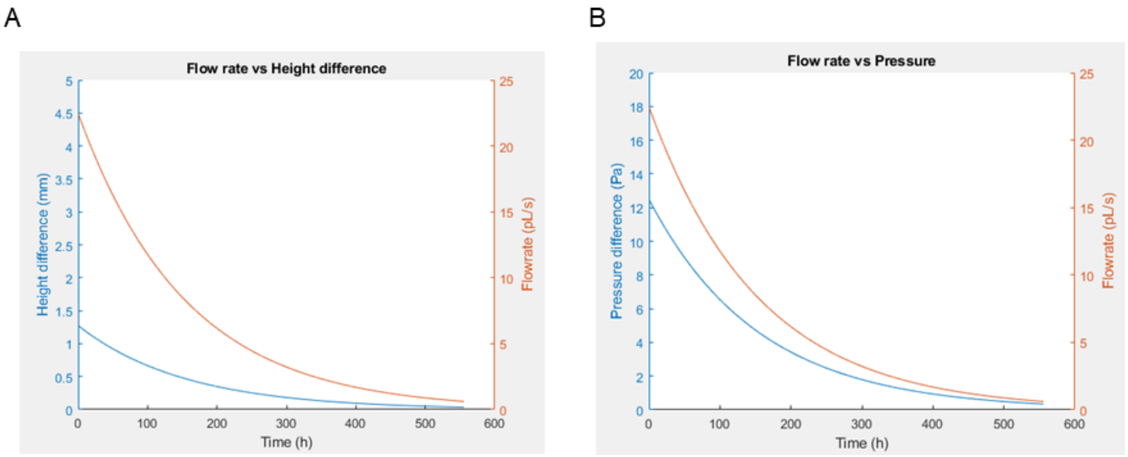


**Figure S10.** MATLAB output graphs for the (A) Flow rate and height difference along the time, and (B) Flow rate and pressure difference along the time.

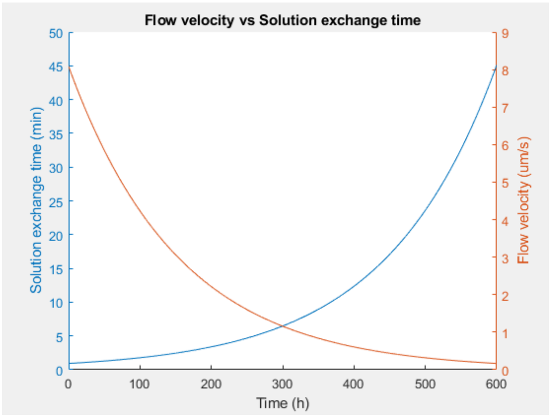


**Figure S11.** MATLAB output graphs for flow velocity and solution exchange time along the time.

#### 6) Axonal transport of neuron-targeted dendriplexes

A

**http://tinyurl.com/TgDendriplexes**

B

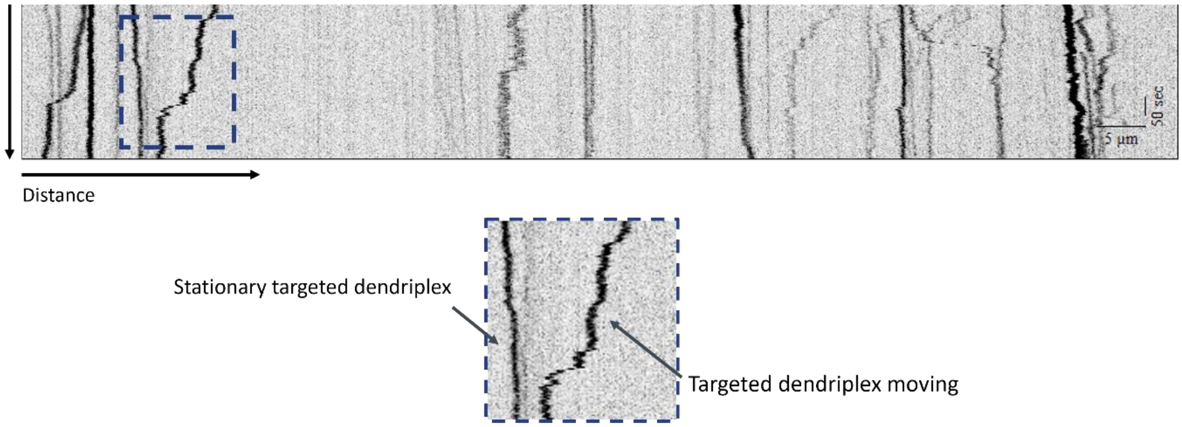


**Figure S11.** A) Representative video of Tg N/P 5 Cy5-siRNAmi-dendriplexes movement in cortical neurons after 18-20 hours of incubation. B) Representative kymograph showing targeted NPs paused or in movement in axons of motor neurons.

#### 7) Evaluation of the capacity to reach cell body of neurons – Axonal compartment

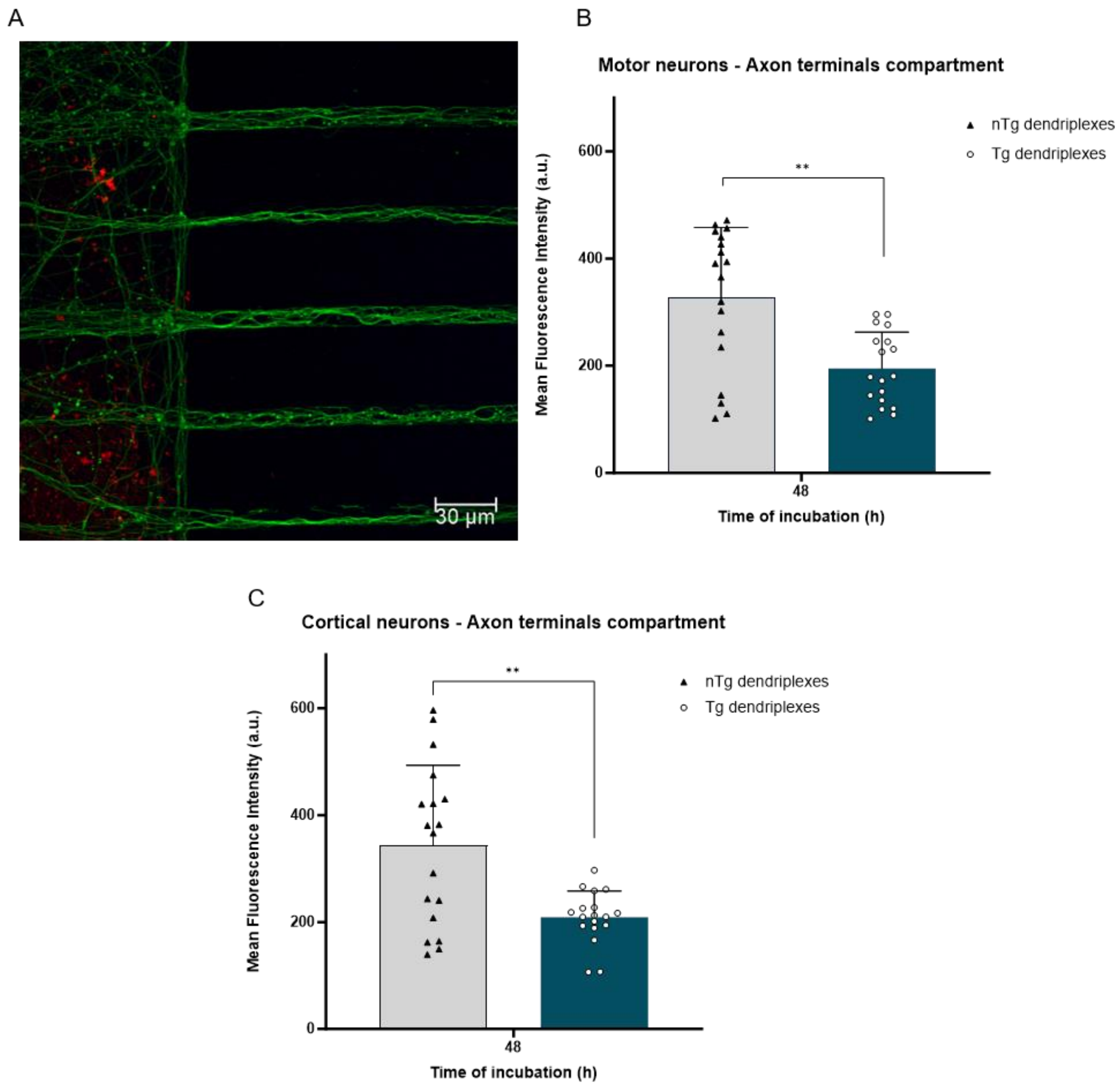


**Figure S12.** Quantification of Cy5 fluorescence inside the axons in the axonal terminals compartment of two-compartment microfluidics. (A) Representative images from confocal microscopy of nTg N/P 5 dendriplexes carrying Cy5-siRNAmi in the axonal terminals compartment of cortical neurons after 48 hours of incubation. Staining: βIII tubulin (in green) and Cy5-siRNAmi dendriplexes (red). Through image analysis, (B) mean Cy5 fluorescence intensity was quantified in the axonal compartment of motor neurons and in the (C) in the axonal compartment of cortical neurons compartment. Results are shown as mean ± SD of three independent experiments (n = 3), 6 replicates per experiment. For statistical analysis, one-way ANOVA test was used. Significant differences: **$p < 0.01$.

#### 8) Biocompatibility evaluation in PNS-CNS-on-Chip

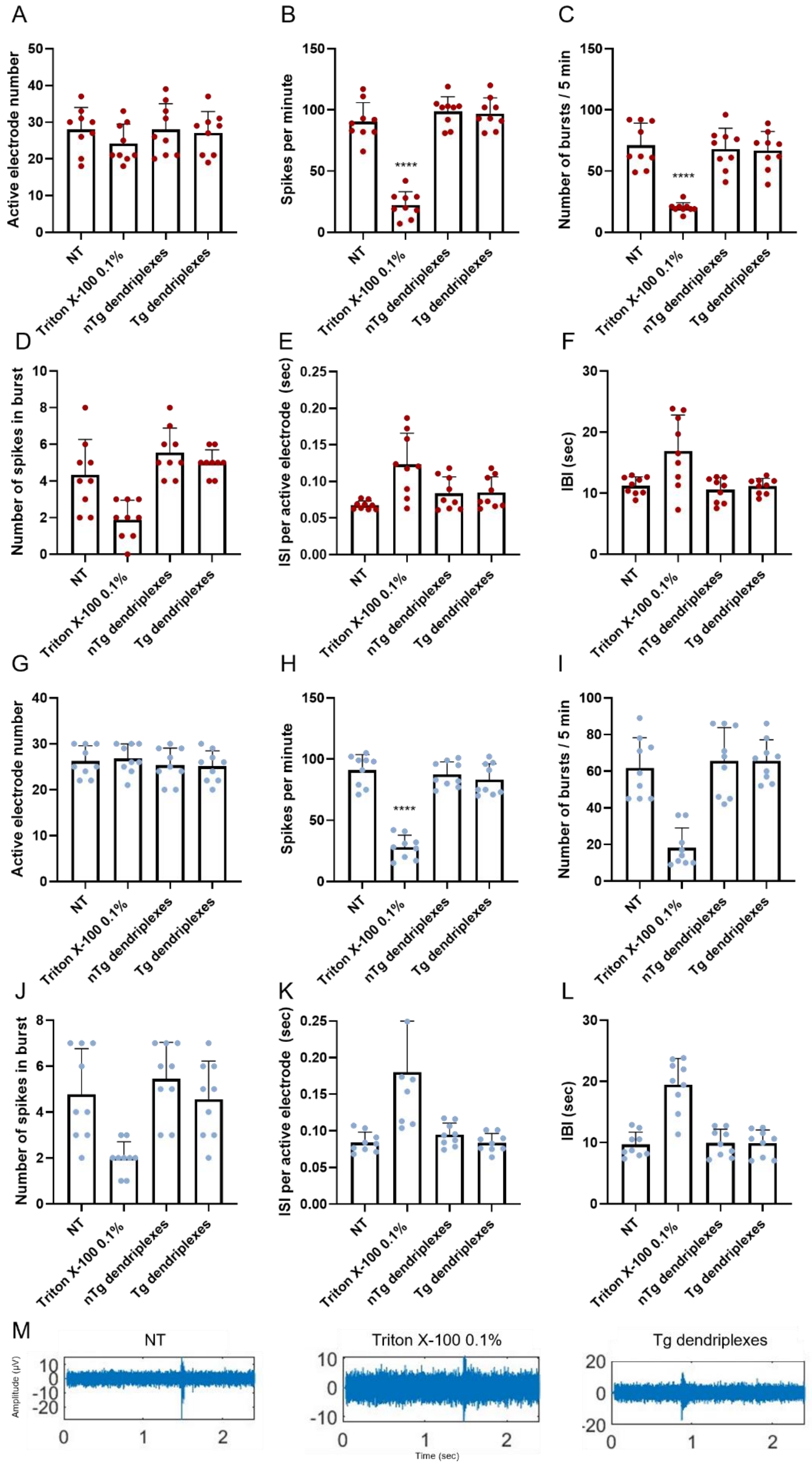

**Figure S13.** Spontaneous electrical activity of (A-F) motor and (G-L) cortical neurons, cultured in PNS-CNS-on-Chip, recorded with microelectrode arrays (MEAs) after incubation with nTg or Tg dendriplexes carrying siPTEN (DIV14). Untreated cells and cells treated with Triton™ X-100, 0.1% (v/v) in PBS 1× were used as controls. (A) and (G) Number of active electrodes. Active electrodes were defined by showing 15 or more spikes per minute; (B) and (H) Number of spikes per minute; (C) and (I) number of bursts during 5-min recording. Bursts were defined as three or more spikes with a 300-ms interspike interval (ISI); (D) and (J) number of spikes in burst; (E) and (K) ISI per active electrode; (F) and (L) mean intra-burst interval (IBI). Results are represented as the mean of three independent experiments (n = 3). For statistical analysis, a two-way ANOVA test was used. No statistically significant differences were identified resulting from treatment with nTg or Tg dendriplexes (N/P 5) compared to the non-treated (NT) cells. A voltage-threshold based algorithm applied to 20,000 Hz high-pass filtered traces was used, with the threshold set to 4.5 times the standard deviation of the noise. (M) Representative spontaneous electrophysiological activity of NT neurons, neurons treated with Triton™ X-100, 0.1% (v/v) in PBS 1×, and neurons treated with Tg dendriplexes (N/P 5). Results are expressed as mean ± SD of three independent experiments (n = 3), with 1 replicate per experiment. For statistical analysis, two-way ANOVA test was used. Significant differences: **** $p \leq 0.0001$.
